# Genome divergence across the Indo-Burman arc: a tale of two peacocks

**DOI:** 10.1101/2022.05.27.493701

**Authors:** Ajinkya Bharatraj Patil, Nagarjun Vijay

## Abstract

Exaggerated traits of the peacock are attributed to sexual selection. Yet, the two species of Asian peacock are conspicuously different in their plumage colouration and level of sexual dichromatism. Our integrative comparative study of bird calls, morphological differences and genomic divergence between the Indian blue peafowl and the green peafowl suggests a strong role of habitat differences in shaping species-specific changes. We estimate a recent (1-3 MYA) split of these species in the Pliocene or early Pleistocene, followed by intermittent periods of gene flow. Despite the relatively recent split, the high levels of genomic differentiation (F_ST_ ∼ 0.6) mirror the divergence in morphological traits. Several genes involved in body patterning and colouration have accumulated protein-coding differences between the peacock species. Our estimates suggest genetic diversity in the widespread Indian peafowl (θ_w_ ∼ 0.0015) is comparable but slightly higher than in the endangered green peafowl (θ_w_ ∼ 0.0012). The ratio of genetic diversity on the Z chromosome to the autosomes (Z/A) is consistent with a polygynous mating system in the Indian peafowl compared to monogamy in the green peafowl. The Asian peacock species continue to provide exciting new insights into speciation and mating system evolution in the post-genomic era.

## Introduction

Diversification of plumage colouration has propelled the radiation of birds (Owens *et al*. 1999). Yet, the role of sexual selection in speciation and divergent plumage colour evolution are hotly debated topics. Macroevolutionary trends in large phylogenetically diverse datasets are used to decipher the relationship between speciation, sexual selection, and plumage colouration (Kraaijeveld *et al*. 2011, Huang & Rabosky 2014, Dale *et al*. 2015, Cooney *et al*. 2019, Price-Waldman *et al*. 2020, Cally *et al*. 2021). The genetic mechanisms of gain/loss of characters between closely related species driving this correlation are understudied. Despite recent speciation, the genus *Pavo* has striking plumage colouration changes, differing levels of sexual dichromatism, and sexual selection (Kimball *et al*. 2011, Jetz *et al*. 2012). Sexual selection is more intense in polygamous than monogamous species (Emlen & Oring 1977). The differing levels of sexual selection between the two *Pavo* species may be a result of a change in the mating system. The Z chromosome to autosome genetic diversity (Z/A) ratio provides a proxy to address a shift in the mating system between closely related species (Corl & Ellegren 2012). Chromosome-level genome assemblies and genomic sampling of multiple individuals make this genus a compelling model for comparative genomics (Jaiswal *et al*. 2018, Dong *et al*. 2021, Zhang *et al*. 2022). For instance, it allows us to compare the Z/A ratio between these species to get insights into mating system evolution.

The two species of the genus *Pavo* (Asian peacocks) and the Congo peacock are the three extant peacock species (Kimball *et al*. 1997). The Asian peacock is an exquisite bird with glamourous colouration and extravagant plumage, which has made it extremely popular even outside its native range (Christine E. Jackson 2006)□. The diverse colours and patterns of its long tail of iridescent feathers, unmistakable crest, loud call pattern, and large body size make them one of a kind. During the breeding season, extreme sexual dimorphism can be seen in peacocks as males develop a long train (∼1-1.2 meters) of feathers with oval multicoloured ocelli (Lin n.d., Christine E. Jackson 2006, Monique de Vrijer 2007)□. The number and patterns of ocelli are decisive in female choice, as they boast about their health and good genes (Petrie & Williams 1993, Møller & Petrie 2002, Loyau *et al*. 2005, Hale *et al*. 2009). The Indian peafowl (*Pavo cristatus*) is more widespread than the green peafowl (*Pavo muticus*) and the more divergent Congo peafowl (*Afropavo congensis*), which are rarer and have a smaller distribution range (Lin n.d., Brickle, N. W., Nguyen Cu, Ha Quy Quynh 1998, Lukanov 2013, Kushawaha & Kumar 2016)□. The Indian peafowl is hardier and more tolerant of human habitations, has a larger population size, and is semi-domesticated in multiple non-native countries worldwide (Lin n.d., Christine E. Jackson 2006, Hernowo *et al*. 2011). The green peafowl is more aggressive and territorial and is quite intolerant to anthropogenic influence, restricting its range to forest areas (Hernowo *et al*. 2011)□.These habitat differences may have played a role in their phenotypic divergence.

The Congo peafowl is the only valid peacock species with a native range outside Asia. It is much smaller than other peacocks and has considerably different morphological traits. Previous estimates of divergence between *Pavo* and *Afropavo* dates approx. 11.4 MYA (Kimball *et al*. 1997)□. The estimates of divergence time between the Indian peafowl (*P. cristatus*) and the green peafowl (*P. muticus*) range from 1 to 6 MYA (Kimball *et al*. 1997, Jetz *et al*. 2012, Kumar *et al*. 2017)□. But these two species, whose current native ranges don’t overlap, still hybridise in captivity as their albumin and transferrin indexes are identical (Prager & Wilson 1975)□. Sexual dimorphism among these species is also an interesting phenomenon. *Afropavo* and *P. cristatus* have a high degree of sexual dimorphism, whereas it seems relaxed in *P. muticus*. The estrogen pathway regulates the maintenance of sexual dimorphism through its downstream targets (Owens & Short 1995, Björnström & Sjöberg 2005)□□. The investigation of the targets of estrogen might prove beneficial in finding underlying genes involved in sexual dimorphism.

Candidate gene-based studies were a popular approach for identifying genetic differences between species until the advent of next-generation sequencing (NGS) technologies. The reduced cost of NGS led to many speciation genomics studies (Seehausen *et al*. 2014). Speciation genomics studies sequenced the genomes of multiple individuals from closely related species and used population genomic metrics like F_ST_ (differentiation) and D_xy_ (divergence) to identify species-specific nucleotide changes (Nosil & Feder 2012, Ravinet *et al*. 2017). Regions of the genome with high differentiation and/or divergence compared to the genomic background are identified as outliers based on various speciation models. Such outlier regions are termed speciation islands and may have a role in speciation. The relevance of these speciation islands and the processes that govern these patterns have been the focus of considerable debate (Wolf & Ellegren 2016). Nonetheless, fixed differences that result in protein-coding changes are likely to have phenotypic consequences. Hence, this approach to finding fixed differences and evaluating their effects is similar to comparative genomics but can distinguish segregating variants from fixed sites.

In this study, we use publicly available genomic and transcriptomic data of the Indian peacock (*P. cristatus*) and genomic data of the green peacock (*P. muticus*) to characterise genome divergence between these species. We also collate various phenotypic differences that could result from the genetic changes. Objectives for this study are:

1. To collate the phenotypic differences between *P. cristatus* and *P. muticus* based on an extensive literature survey.
2. To perform acoustic analysis of their calls to identify distinguishing features.
3. To estimate the divergence time between the two species using two complementary methods, (a) UCE-based phylogeny and (b) PSMC pseudo-diploid analysis.
4. Robustly estimate genetic diversity and use the Z/A gene diversity ratio to assess the mating system change.
5. To find key genetic differences between these two species that might play a role in phenotypic divergence and speciation.

We hypothesise that the striking phenotypic differences between the Asian peacock species result from nucleotide sequence differentiation in various body patterning and colouration pathways. We use a population genetic approach to identify highly differentiated genes between the two species of peacocks. In contrast to a comparative genomic approach, which relies on single sequenced individuals, we can distinguish segregating polymorphisms from strongly differentiated alleles, which may be fixed between the two species. In our dataset, > 50% of (∼5500) of the annotated protein-coding genes are covered by >= ten reads in each of the 12 individuals consisting of six *P. cristatus* and six *P. muticus* individuals. The stringent criteria used in our study resulted in partial coverage of the exome; however, the fixed differences identified are well supported by the data. We find a prevalence of protein-coding fixed differences in several promising candidate gene loci from body patterning and colouration pathways. We propose that the morphological differences in the peacock are a conglomerate of multiple development-related patterning genes that acts differently in these two species.

## Materials and Methods

(See Electronic Supplementary Material for detailed methods)

### (a) Genome annotation

We downloaded the *P. muticus* genome (Dong *et al*. 2021) and the associated SRA reads from NCBI (**Table S1**). The genome was annotated using the Maker 3 (Campbell *et al*. 2014) annotation pipeline. The transcriptomic data from *P. cristatus* was assembled using Trinity version 2.13.2 (Haas *et al*. 2013)□. The obtained transcriptome assembly was used as EST evidence for the annotation. We used the annotations to obtain the candidate genes and verified the ORF using the *Gallus gallus* (galgal7) genome to overcome the annotation errors.

### (b) Acoustic Analyses

The representative calls for *P. cristatus* and *P. muticus* were retrieved from the Xeno-Canto animal call database (Xeno-canto n.d.)□. Calls with minimal background noise (*P. cristatus* -23 and *P. muticus* -20) from 10 recordings (5 from each species) were selected for the analysis in Raven pro 1.6.1 (Yang 2021). The maximum energy bands from the spectrogram were identified as Dominant frequencies. The acoustic measurements (call duration and frequencies) obtained from Raven were analysed in R (R Core Team 2021)□. (see **Table S2** and **S3**)

### (c) Gender identification

We used Bedtools to calculate the number of reads in 50 kb windows for each bam file. To identify gender, we estimated the normalised read counts (i.e., number of reads in a 50 kb window/ mean of the number of reads in all 50 kb windows) of autosomes, Z and W chromosomes for each individual (**Figure S1-S10**). Based on a molecular assay, *P. cristatus* individual-1 (SAMN05660020) gender is known to be male. Keeping this individual as a reference, we divided the normalised read counts in each 50 kb window of *P. cristatus* individual-1 by the normalised read counts in the same 50 kb window in each of the other individuals. Using these two approaches, we identified the genders of previously unidentified individuals (see **Table S1**).

### (d) Population genomic summary statistics

We mapped sequencing reads of 6 samples each from *P. cristatus* and *P. muticus* to the genome sequence of *P. Muticus* (Zhang *et al*. 2022). The 2D-SFS between *P. cristatus* and *P. muticus* was used to generate per base, 1KB and 50 KB non-sliding window F_ST_ using the FST stats2 module of ANGSD (Korneliussen *et al*. 2014)□. The SAF files were used for summary statistics calculation to get theta estimates using realSFS saf2theta and thetaStat modules for window sizes of 1 KB and 50 KB non-overlapping windows. We calculated the mean estimates of population genomic statistics such as Watterson’s theta (θ_w_), Pairwise theta (θ_π_), Tajima’s D (τ) and F_ST_ for autosomes and the Z chromosome separately. We estimated these statistics in 1 KB windows with varying levels (at least 400 to 900 sites covered) of missing data to ensure that data quality does not affect our estimates (**Figure S11-S20**). We calculated the Z/A ratios of all individuals and males only separately using the corresponding diversity estimates.

### (e) Shortlisting of the candidate genes

Fixed sites (F_ST_ > 0.9) identified by ANGSD were intersected with the vcf file generated by bcftools using Bedtools (Quinlan & Hall 2010)□ to get a fixed-sites vcf file. We used the fixed-sites vcf file to make an alternate fasta sequence for each species using AlternateFastaMaker of GATK v 4.2.0 (Van der Auwera *et al*. 2013).

We employed two approaches to identify candidate genes. The first approach annotated the fixed-sites vcf file using SnpEff (Cingolani *et al*. 2012) and prioritised the candidate genes based on the gene-specific ratio of non-synonymous to synonymous changes. The exons were prioritised based on a) the ratio of fixed differences upon their length (i.e., fixed-site density) (101 of 200 highly differentiated genes have >= one amino acid difference). (b) All genes on the Z chromosome (95 of 887 genes had >= one amino acid difference). (c) previously reported melanogenesis-related genes (20 of 145 genes had >= one amino acid difference) (see **Table S4**).

As part of the second approach, genes involved in KEGG signalling pathways (Kanehisa *et al*. 2017)□ and other related genes were considered candidates. The corresponding protein sequence from *G. gallus* (galgal7) was used for each candidate gene to query *P. cristatus* and *P. muticus* genomes using Exonerate (Slater & Birney 2005) protein2genome tool. We compared the translated coding sequences to identify differences between the peacock species (see **Table S5**).

### (f) Demographic history

The genomic bam alignments for each species were used to get consensus calls using samtools (Li *et al*. 2009) and bcftools (Li & Barrett 2011)□. Obtained fastq calls were converted to the psmcfa file using the fq2psmcfa command of the psmc suite (Li & Durbin 2011)□. The resultant psmcfa file was then used to execute psmc with options -N30 -r5 -t5 -p 4+30*2+4+6+10. The estimated trajectory of effective population size (N_e_) was plotted with a mutation rate (u) of 1.33e-09 per site per year (Wright *et al*. 2015)□ with a generation time of four years (Tacutu *et al*. 2018) using psmc_plot.pl script of psmc.

### (g) Divergence time estimate

We employed two independent methods to obtain divergence time estimates. Genomic UCE-based alignments and pseudo-diploid analysis of psmc were performed to get estimates of divergence between these two species. We followed the Phyluce-1.7.1 (Faircloth 2016) pipeline to get concatenated alignments of 75% of representative gene trees. After obtaining bootstrapped trees from raxml-ng (Kozlov *et al*. 2019), MCMCTree from PAMLv4.9f (Yang 2007)□ was used to estimate the divergence time.

The WGS reads from both species were mapped to the *P. muticus* genome. The haploid genomes from both species were merged to produce a pseudo-diploid genome using Seqtk (Li 2013). The parameters (-N25 -t5 -r5 -p “20+4*5”) were used for psmc. The time at which the effective population size (N_e_) goes to infinity was considered the divergence estimate for those two species.

## Results and Discussion

### (a) Plumage and acoustics are strikingly different

The plumage colouration is the most striking difference between the Indian blue peafowl and the green peafowl. Although the plumage colour is different, both species have long train feathers spaning 1-1.5 meters right after the backplate. The train feathers are around 130-150 in number, with greenish-blue ocelli at the end and about 20-30 V-shaped apical feathers (Lin n.d., Christine E. Jackson 2006, Lukanov 2013, Kushawaha & Kumar 2016). The peacock’s true tail consists of ∼20 powerful tail feathers, which are instrumental in raising the train feathers. The wing feathers help shimmer the whole train during the display **(Figure 1A (a)** and **(d)**). The green peafowl is slightly larger than the blue peafowl despite weighing less. The adult male of *P. cristatus* weighs around 4-6 Kgs and spans around 2-2.3 meters, whereas the female is slightly lighter in weight (2-4Kgs) and smaller (about 1 meter) due to the absence of train feathers. The adult male of *P. muticus* is comparatively lighter (4-5 Kgs) but has a larger body size of 2.5-3 meters, and females are just longer than 1.1 meters (Lin n.d., Christine E. Jackson 2006, Lukanov 2013, Kushawaha & Kumar 2016).

**Figure 1A.**
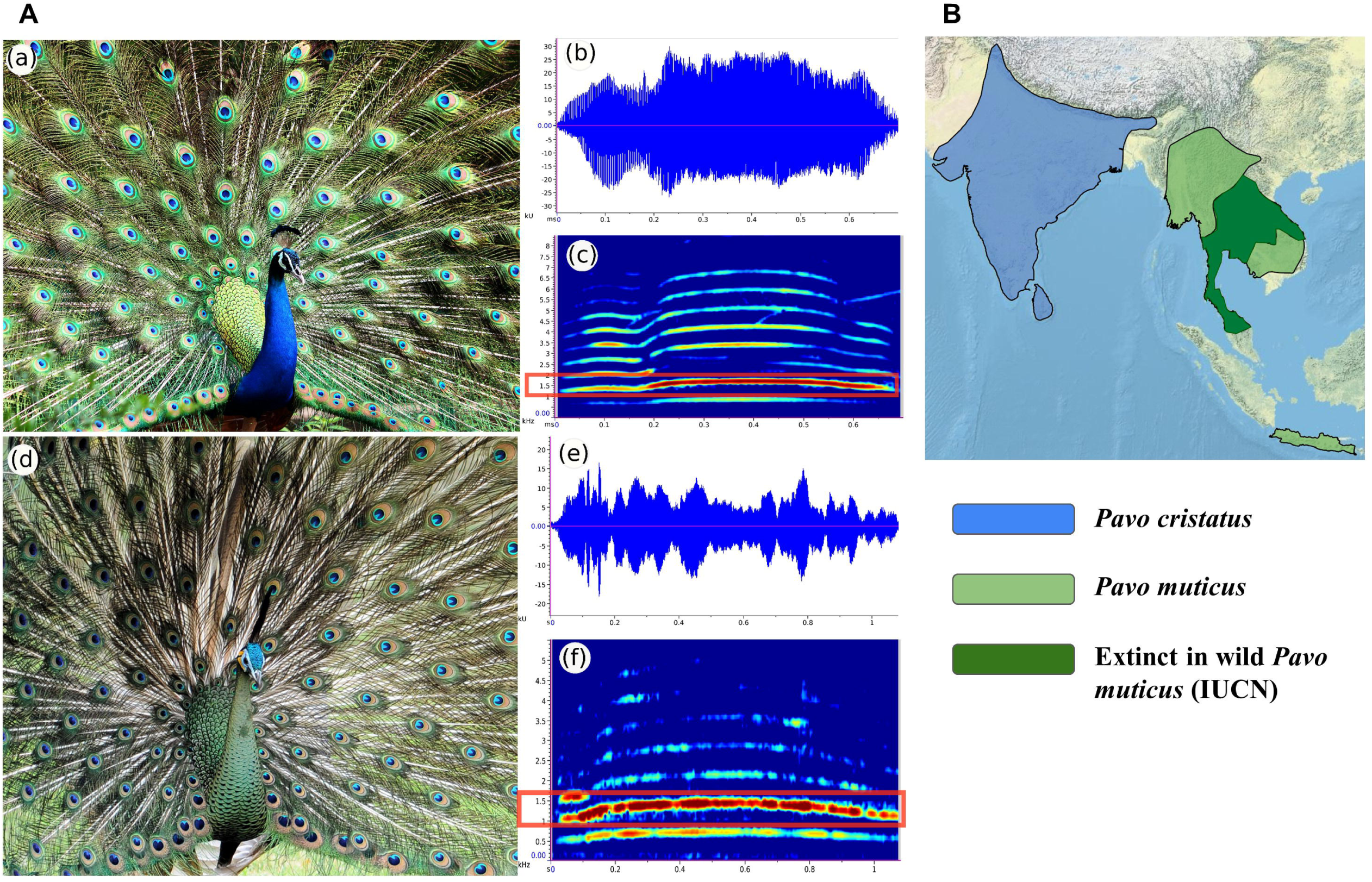
Acoustic differences between the Indian blue peafowl (*Pavo cristatus*) and the green peafowl (*Pavo muticus*). (a) The male of Indian blue peafowl (*P. cristatus*) and (b) The waveform view of a single call of Indian blue peafowl (*P. cristatus*). The call lasts for approximately 0.7 seconds. (c) The spectrogram view of a single call of Indian blue peafowl (*P. cristatus*). The call comprises multiple harmonics with its fundamental frequency starting at ∼800 Hz and second harmonics is the dominant frequency at ∼1600 Hz. (d) The male of the green peafowl (*P. muticus*) during its sexual display. (e) The waveform view of a single call of the green peafowl (*P. muticus*). The call lasts for approximately 1 second. (f) The spectrogram view of a single call of the green peafowl (*P. muticus*). The call is composed of multiple harmonics, with its fundamental frequency starting at ∼600 Hz and second harmonics being the dominant frequency at ∼1200 Hz. **Figure 1B: The geographic native distribution of *Pavo cristatus* and *Pavo muticus***. The Indian blue peafowl is distributed throughout the Indian subcontinent, including India, Sri Lanka, Pakistan, Bangladesh, Nepal, etc. (highlighted in blue). The green peafowl has a relatively smaller distribution in Myanmar, Cambodia, Thailand, Vietnam, Indonesia, etc. (highlighted in green). According to the IUCN database, the species might be extinct in the wild in and around Thailand (highlighted in dark green).

The call of peacocks is considered one of the loudest among birds. Both the species of peacocks have a call with multiple harmonics with second harmonics as their dominant frequency, as seen on the spectrogram (**Figure 1A(c)** and **(f)**). *P. cristatus* (∼ 0.653 seconds, σ = 0.072) has a significantly shorter, less variable call than *P. muticus* (∼ 0.908 seconds, σ = 0.195) **(Figure 1A (b), (e), S21-S24** and **Table S2)**. The harmonic structure looks similar between the two species, but they differ in their maximum frequencies among their harmonic bands. The fundamental frequency of *Pavo cristatus* (∼756 Hz) is significantly higher than *P. muticus* (∼551 Hz) (**Figure S24**, Pairwise Wilcoxon test, p < 0.0001). The second harmonics have the highest energy band (dominant frequency) in both the species, with *P. cristatus* (∼1489 Hz) having a significantly higher maximum frequency than *P. muticus* (∼1111 Hz) **(Table 2, Figure 1A, S22, and S23)** (Pairwise Wilcoxon test, p < 0.0001). Hence, the temporal and spectral call properties are distinctive, illustrating the different calls among the two species of peafowls. Species’ habitat shapes bird song through changes in body size which is an important determinant of bird call properties. Lowland forest species have low frequency and long calls due to the larger body size (Ryan & Brenowitz 2015, Marcolin *et al*. 2022). The lower-frequency, longer calls of *P. muticus* are likely to be adaptations to the lowland forest habitat and may be responsible for their higher susceptibility to anthropogenic change (Dong *et al*. 2021).

The Indian peafowl (*P. cristatus*) is distributed throughout India, and its native range covers the elevation gradient of lowland deciduous forests to the peaks of Western Ghats. According to the IUCN red list database, The green peafowl is sparsely distributed in the Southeast Asian region from Myanmar to Laos, Cambodia, Thailand, Vietnam, Malaysia, and Java. The two species seem to be separated by the Indo-Myanmar mountain range, which continues to the Himalayas (**Figure 1B**). The current geographic distribution of these two species can be traced back to dispersal events in the late Miocene to Pliocene following major geological changes (Ripley & Beehler 1990).

With a restricted geographic range and greater habitat specificity, the *P. muticus* populations are smaller and fragmented, classifying them as endangered species. With a broader geographic range *P. cristatus* although classified as a species of least concern, enjoys legal protection throughout most of its range, being the National bird of India. Our estimates suggest that the nucleotide diversity of *P. muticus* (θ_w_ =0.00124, θ_π_ =0.00116) is lower than that of *P. cristatus* (θ_w_ = 0.00149, θ_π_ = 0.00151) (see **Table S5**). Although our estimates of the genetic diversity of *P. muticus* are consistent with previous reports, more geographically widespread genome-wide sampling will be able to provide robust measures of the diversity in both species.

### (b) Colouration, body, and feather patterning have diverged

The male peacock is an ornamental stock with multiple types of feathers with various colours around the body. While the *P. cristatus* has a crest with apical iridescent blue feathers which stand on the black stalk, arranged in a thin-striped fan-shaped manner, the *P. muticus* has a long, dense, tufted crest with bluish-green colour to its feathers. The crest is leaning forward on the head in the green peafowl and is located somewhat further than the blue peacock in which the crest is slightly leaning backwards (**Figure 2A-A**). The facial area of *P. cristatus* extends from its beak to the ear through crescent-shaped white featherless skin, with small bluish-green head feathers. The blue feathers of the neck continue till the back, where the feather pattern changes to a greenish scaly backplate, which appears as primordial trail feathers with smaller ocelli in the centre. The *P. muticus* has a larger featherless area on its face, with black colour in the frontal area of the eye, and blue-tinged white crescent-shaped skin, surrounded by yellow featherless skin on its cheek area. Its head and neck feathers have a scaly pattern similar to backplate feathers and are metallic green (**Figure 2A-B to E**). The top wing feathers of *P*.*cristatus* are primarily brownish with black stripes and a bluish-black bottom. The bottom feathers are reddish, whereas other belly feathers have a brownish-black tinge. The wing feathers of *P. muticus* are utterly different from Indian peacocks, as they are dark black coloured with bluish feathers at the top with cream-coloured bottom feathers (**Figure 2A-F and G**). Although the trail feathers found in the males are breeding season-specific, the other morphological features are consistent throughout the year. Hence, the between-species differences in feather patterning and colouration are likely caused by genetic changes in cell migration and differentiation pathways.

**Figure 2.**
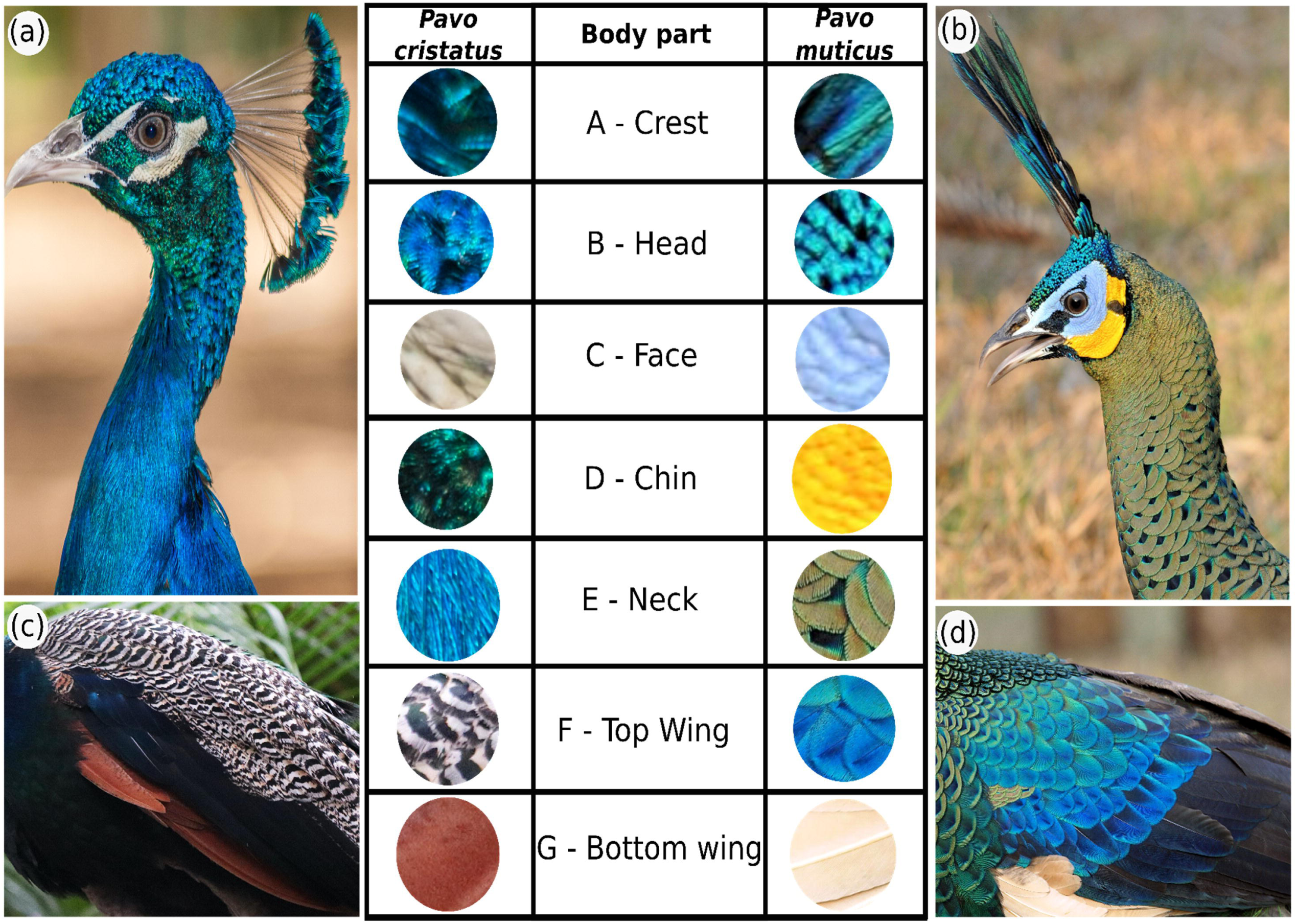
The morphological differences between the male Indian blue peafowl (*Pavo cristatus*) and the male green peafowl (*Pavo muticus*). A. The Indian blue peafowl neck and (c) body feathers. (b) The green peafowl neck and (d) body feathers. The crest of the peafowls. The Indian blue peacock has a thin strip of the fan-shaped crest at the back of its head with iridescent blue feathers at its apex. The green peafowl has long tufted blackish-blue feathers as a crest at the central part of its head pointing up and forward. B. The head of the peafowls. The head of the Indian blue peacock is rounded with short bluish feathers, whereas the green peafowl has a downward slanted head with short green-bluish feathers. C. The face of the peafowls. The face of the Indian blue peacock has pale, whitish featherless skin surrounding the eye, which is interrupted by a small patch of greenish feathers in front of the eye. The green peafowl has a greyish white patch covering the eye, interrupted by a small patch of blackish-green featherless skin. D.The chin of the peafowl: The chin of the Indian blue peacock has greenish feathers extending up to the neck, whereas the green peafowl has a yellow featherless patch with black borders which does not extend to the neck. E.The neck of the peafowl: The Indian blue peacock has an even distribution of blue, iridescent, barbule-like small feathers all over its neck until its back. The green peafowl has scale-like feathers that appear greenish when stacked close together, whereas the distal part of the feather has green and bluish-black spots, which resemble the immature ocelli feathers pretty closely. F.The top feather of peafowls. The Indian blue peacock has barred feathers with creamy-brown colour with intermittent black stripes. The green peafowl has two colour gradients of the top feather. The initial part starts with bright bluish feathers with dark black colour on the other half. G.The bottom feathers of peafowls. The Indian blue peacock has reddish-brown coloured feathers, whereas the green peafowl has creamy coloured feathers.

### (c) Mating system evolution

The most glaring difference between green and blue peacocks is their females. The peahen of *P. cristatus* is dull with no train and whitish-brown body feathers, with a minor patch of greenish scaly feathers around the neck (**Figure S25**). The *P. muticus* females are pretty similar to the males, with identical body feathers and colouration. Their sexes are tricky to identify outside the breeding period **(Figure S26)**. The *P. cristatus* peahen is comparatively more fecund and can lay around 5-8 eggs at once and about 28 eggs in a season in captivity. The *P. muticus* females lay about 3-6 eggs once and 10-12 eggs a season. The blue peacocks are known for breeding in larger groups or “leks”, and the breeding system looks polygynous, with males courting multiple females with no or negligible parental care. Considering the similar morphologies of males and females, the green peacocks seem to have a different breeding system than their Indian counterparts. Their call is comparatively milder than Indian male peacock. The male is highly territorial, defending their area aggressively, and shows parental care suggesting a monogamous mating system. The ratio of genetic diversity of the sex chromosome to the autosomes (Z/A ratio) is different between monogamous and polygamous species (Ellegren & Galtier 2016).

In polygynous species, the Z/A ratio is slightly less than that of monogamous, potentially due to the effects of sexual selection (Corl & Ellegren 2012). We found the Z/A ratio in *P. cristatus* (0.45) is less than in *P. muticus* (0.58), suggesting that *P. cristatus* is polygynous in contrast to monogamous *P. muticus* (see **Table S5** and **S6**). Other than the mating system, demographic history, life history, domestication, etc., can affect the Z/A ratio. Since the two Asian peacock species are closely related, most of these other factors are comparable. However, the demographic histories are different despite the declining population size in both species. Our estimate of Tajima’s D for *P. muticus* (τ = -0.23) may reflect the prevalence of admixed populations in *P. muticus* (excess of low-frequency mutations). The positive values of Tajima’s D in *P. cristatus* (τ = 0.04) may either be due to recent population expansion (excess of high-frequency mutation) or an artefact of limited sampling. Nonetheless, the differences in the Tajima’s D estimates don’t appear to be strong enough to explain the Z/A ratio differences between the two Asian peacock species.

### (d) Peafowl species have diverged from the late Pliocene to the early Pleistocene

The divergence time of *P. cristatus* and *P. muticus* is ∼2 MYA according to the UCE-based dated phylogeny (**Figure 3A**). The previously published divergence time estimates for this species pair range from 1 MYA to 6 MYA (**Figure 3B**). Excluding one mitochondrial marker-based study (Kimball *et al*. 1997) narrows the divergence range from 1 to 3 MYA. The pseudo-diploid analysis suggests a split time of ∼3 MYA, followed by gene flow between 1.8 MYA and 1.1 MYA (**Figure 3C and Table S7**). Overall the UCE-based estimate, previously published studies, and pseudo-diploid analysis support a divergence time between 1.1 MYA to 3 MYA. The heterogeneity between divergence time estimates from various studies decreases with increasing data and supports a much more recent split time with periods of gene flow. Previous biogeographic studies have claimed that speciation in the Asian peacock is mainly due to dispersal (Ripley & Beehler 1990). However, we find that intermittent periods of gene flow due to glaciation events occurred after an initial split due to dispersal in the late Pliocene. During glaciation, the colder climates favoured gene flow due to increased connectivity between the two species. The warmer temperatures during the interglacials resulted in altered species distributions, reducing connectivity between the two species (Dong *et al*. 2021). Hence, dispersal and glacial cycles both contributed to the speciation of Asian peafowls. Similar speciation patterns in the Pliocene to early Pleistocene are reported in the closely related peacock pheasants (*Polypectron* genus), which partly share the geographic distribution with the Asian peacock (Kimball *et al*. 2001).

**Figure 3C.**
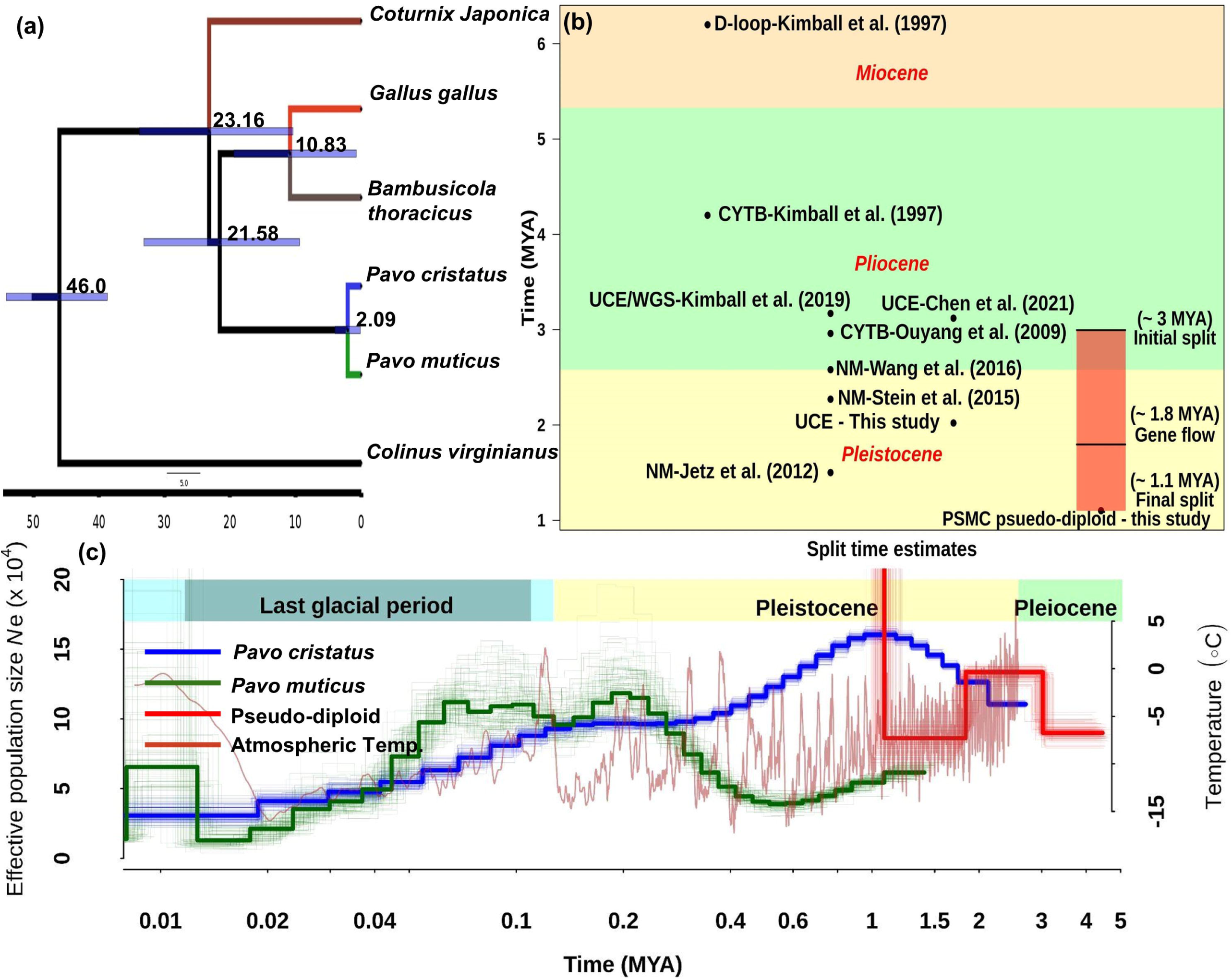
The demographic history and divergence time estimates between *Pavo cristatus* and *Pavo muticus*. (a) Whole-genome UCE-based divergence time phylogeny estimates approximate split times at the nodes. *Colinus virginianus* is used as an outgroup, and this root is placed at 46 MYA. The blue rectangles denote 95% confidence intervals (CI) for these divergence estimates. The split time between *P. cristatus* and *P. muticus* is estimated at ∼2 MYA. (b) The compilation of divergence time estimates between *P. cristatus* and *P. muticus* from the current study and previously published literature. The type of data used in the respective studies is denoted by the short prefixes such as CYTB - Cytochrome B, NM - Multiple nuclear and mitochondrial markers, UCE - Ultra-Conserved elements, WGS - whole-genome sequencing, etc. Most of the divergence time estimates lie in between 1 to 3 MYA. The red rectangle denotes the PSMC pseudo-diploid-based scenario of the divergence between these two peafowl species. The first split is around 3 MYA during the late-Pleiocene and the early-Pleistocene transition. From ∼1.8 to 1.1 MYA, there seems to be a gene flow between these two species. Finally, there is a subsequent split at ∼ 1.1 MYA. (c) The demographic history of *P. cristatus* suggests expansion in population sizes till 1 MYA, after which it undergoes contraction till 200 KYA. After brief stability, the population sizes again decreased since the onset of the last glacial period. *P. muticus* also suggests a decrease in population sizes since 1 MYA, but their numbers have increased from 580 KYA to 200 KYA. After a brief upheaval in numbers, the contraction starts around 60 KYA drastically. The red curve denotes the pseudo-diploid estimates between these two species. The curve suggests first split around 3 MYA, maintained till 1.8 MYA. From 1.8 MYA to 1.1 MYA, there was a period of gene flow between these two species, followed by a final split at ∼1.1 MYA. Colder climatic conditions during the late-Pleiocene to the early-Pleistocene transition might have played an instrumental role in the divergence of these two species.

*Pavo cristatus* population sizes increased during the early Pleistocene till 1 MYA. The demographic analysis was unable to reconstruct the history of *P. muticus* prior to 1.5 MYA, but it may have already begun to decline. The population sizes of both *P. cristatus* and *P. muticus* declined after 1 MYA till 500 KYA. However, after 500 KYA, the population size of *P. muticus* began to increase while *P. cristatus* kept decreasing till the start of an interglacial period (∼250 KYA) when it stabilised. *P. cristatus* population sizes were stable till 110 KYA when it began a continuous decline. The increase in population sizes of *P. muticus* continued from 500 KYA to 190 KYA. During the last glacial period, the *P. muticus* population size increased slightly between 110 KYA till 50 KYA before beginning to decline. While the population sizes of *P. cristatus* and *P. muticus* have declined from 50 KYA, the decline has been more pronounced for *P. muticus* than for its Indian counterpart. The recent decline in the population size of *P. muticus* is due to anthropogenic effects (Dong *et al*. 2021).

### (e) Patterning pathway differentiation mirrors the striking phenotypic differences

We found at least one amino acid difference in 50 of 801 genes from 9 selected KEGG pathways related to the differing morphological traits between these species (**Table S8 and Figure S27-S34**). The most number of differences occur in three genes, with 4, 5, and 6 amino acid differences in *ZFYVE16* or Endofin (TGF-Beta), *APC* (Wnt and Actin cytoskeleton), and *LAMA5* (ECM interaction and Focal adhesion) respectively. Endofin (*ZFYVE16*) is a regulator of the TGF-Beta pathway, which regulates positive and negative feedback of the BMP-SMAD signalling. *APC* is a regulator of the Wnt canonical pathway, which acts as an antagonist by downregulating β-catenin (*CTNNB1*) and plays a vital role in microtubule stabilisation in the Actin cytoskeleton pathway. *LAMA5* is one of the essential glycoproteins of the basement membrane and a part of extracellular matrix interactions. *LAMA5* is also a crucial player in cell migration, differentiation, adhesion, etc.

Along with *APC*, four other genes of the Wnt-signalling pathway, namely *GPC4, INVS, PLCB1*, and *ROR2*, have amino acid differences. *GPC4, INVS*, and *ROR2* are involved in the Planar Cell Polarity (PCP) pathway, which determines the cell polarity across the tissue plane and is thus instrumental in patterning. At the same time, *PLCB1* works in the Wnt/Ca^2+^ pathway and is known to be involved in Melanogenesis (**Figure 4**). Genes from multiple patterning signalling pathways such as Hedgehog, Notch, TGF-Beta, and Melanogenesis have peptide sequence differences between the two species (see **Supplementary Notes and Figure S35**). The consequences of these amino acid changes in several genes of the patterning pathway will require experimental evaluation. Nonetheless, the candidate genes identified by this initial screen provide clues about the importance of the divergence in key genes of the patterning pathway. The relatively high differentiation (F_ST_ ∼ 0.6) between the two species makes it challenging to identify signatures of selective sweeps in these species. However, extensive sampling of peafowl populations spanning the entire geographic distribution and understanding the population structure within each species may help disentangle the species-specific differences from population-specific ones.

**Figure 4.**
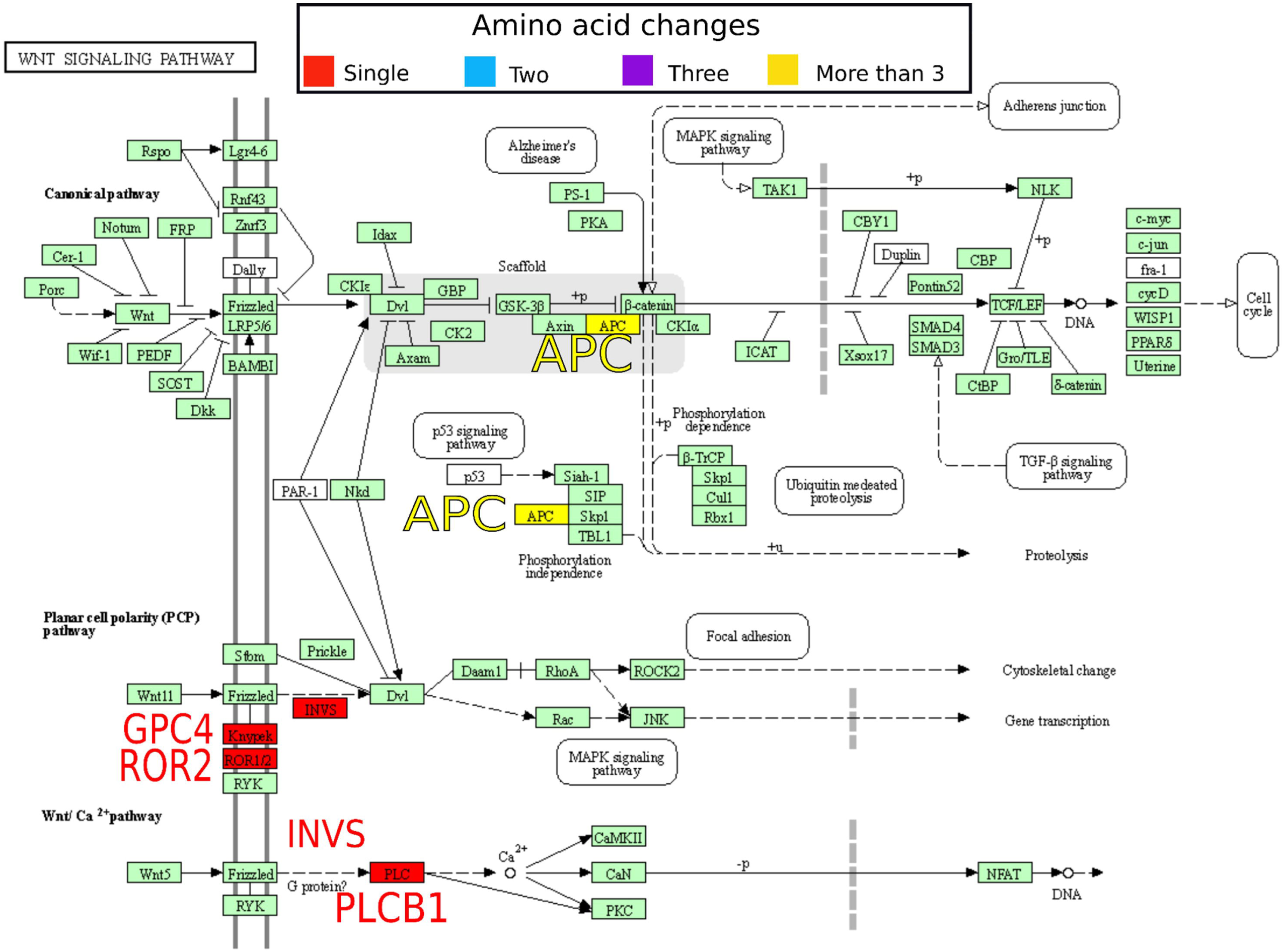
The KEGG annotation of the Wnt signalling pathway. Five genes have amino acid differences between the peafowl species. GPC4, ROR2, INVS, and PLCB1 have a single amino acid difference (highlighted in red between the two species, whereas APC (The Wnt pathway regulator) has a total of 5 amino acid differences (highlighted in yellow).

### (f) The extent of sexual dimorphism is diminished in *Pavo muticus*

*P. cristatus* has extreme sexual dimorphism, also observed outside the breeding season (**Figure S25**). The male has soft bluish feathers throughout the neck, whereas the female lack these feathers. The female has scaly colourless feathers with iridescent greenish colour at the top of her neck. The male has different colours for different body parts throughout the body, whereas the female has white feathers with a brownish-black tinge. In contrast, *P. muticus* does not have any morphological differences except in the male’s tail feathers during the breeding season (**Figure S26**).

The maintenance of sexual dimorphism is primarily attributed to estrogen and its downstream regulation. The MAPK pathway has been the prime target of the estrogens and plays a vital role in its regulation. We found 15 genes with amino acid differences between the two peacocks (see **Table S8** and **Figure S34**). Among these genes, CREB domain-containing *ATF4, IGF2, INSR, NF1, CHUK*, and *NFKB2* seem to be the most critical targets among many other genes of the pathway with sequence divergence. In addition to the fixed differences seen within the protein-coding regions, changes in cis-acting regions would also influence the regulation of these pathways. A study of the transcriptional level differences between these species and the genetic basis of expression differences will help uncover the contribution of regulatory changes.

Despite the striking divergence in plumage, acoustics and behaviour, these two species can still hybridise (Prager & Wilson 1975). Nonetheless, these two species’ demographic trajectories are different due to their differential response to glacial cycles. The green peafowl occurs in a lowland forest habitat in contrast to the widespread distribution of the blue peafowl in diverse habitats. This difference in the habitats may be responsible for the differences in acoustic characteristics, body size, and sexual dimorphism due to a change in the intensity of sexual selection following a change in the mating system. The change in plumage colouration may also be a consequence of habitat-specific changes.

## Conclusion

Our study offers insight into the speciation of Asian peacock species based on a comparative analysis of feather patterning, plumage colouration, bird calls and genome sequencing datasets. We conducted an extensive literature search and qualitatively identified the morphological differences between *P. muticus* and *P. cristatus*. Our acoustic analysis of peacock calls indicates that the forest-dwelling *P. muticus* has lower-frequency, longer calls than *P. cristatus* which might be habitat-specific adaptations. The UCE based phylogenetic tree and whole-genome PSMC pseudo-diploid analysis suggest a split time between 1 to 3 MYA with intermittent gene flow between the two Asian peacock species. Reconstruction of the demographic history suggests a strong role of glacial cycles and dispersal in their speciation. Despite the recent divergence and prevalence of gene flow, we estimate a relatively high level of inter-species genetic differentiation (F_ST_ ∼ 0.6) based on currently available sampling. The striking morphological differences and rapid speciation in peacocks may be a general trend in the sub-continental residents of the Pavoninae subfamily. The Z/A ratio of genetic diversity is consistent with a monogamous mating system in *P. muticus* (0.58) and a polygynous mating system in *P. cristatus* (0.45). Using stringent criteria for identifying fixed differences between the species, we find several candidate genes in the patterning pathway (Wnt, BMP-SMAD and MAPK) that may help explain the changes in feather patterning, sexual dimorphism and colouration. Our findings provide a framework for studying speciation in this colourful Asian bird system with conservation concerns.

## Supporting information

Electronic Supplementary Materials

## Author contributions

A.B.P designed the study and collected and analysed the data. A.B.P. wrote the manuscript with inputs from N.V.

## Data accessibility statement

All data and scripts are available on the github repository: https://github.com/Ajinkya-IISERB/Pavo.

## Acknowledgements

We thank the Ministry of Human Resource Development fellowship to ABP. The Department of Biotechnology, Ministry of Science and Technology, India (Grant no. BT/11/IYBA/2018/03) and Science and Engineering Research Board (Grant no. ECR/2017/001430) provided funds used to purchase computational resources (i.e., Har Gobind Khorana Computational Biology cluster) used. We thank Friedrich Esser for his kind gesture in providing us with the pictures of peafowls. We thank Dr. Anand Krishnan for his useful suggestions and discussion.

