## Supplementary material for "Genome divergence across the Indo-Burman arc: a tale of two peacocks": Electronic Supplementary Materials

**Electronic supplementary information**

### Supplementary Notes

#### Patterning pathway differentiation mirrors the striking phenotypic differences

*KIF7*, which has two amino acid differences, is a key regulator of the hedgehog pathway and an important factor for epidermal differentiation. *EFCAB7*, *HHAT*, and *MGRN1* have one amino acid difference (**Figure S28**). Each of these genes is involved in regulating epidermal differentiation or trafficking processes. Both the Delta ligand (*DLL1* and *DLL4*) genes of the Notch pathway have one amino acid difference (**Figure S30**). Apart from Endofin (*ZFYVE16*), a key regulator of the TGF-beta pathway, *RBL1* and *THBS1* have two and one amino acid differences, respectively. *RBL1* is a vital gene regulating cell cycle and division, controlling chromatin through heterochromatin organisation. *THBS1* is involved in cell to cell/matrix interactions and helps form and maintain multiple epidermal structures (**Figure S27**). All these pathways, including Wnt, are involved in embryonic development and morphogenesis, which shows a conglomerate of changes in these two species showing differentiation. *PLCB1* and *CREB3* are the essential genes that showed differences in the Melanogenesis pathway. *CREB3* is a critical transcription factor that, upon phosphorylation, activates *MITF* and generates downstream enzymes to produce melanin. *PLCB1* catalyses the *DAG* (Diacylglycerol) and *IP3* (Inositol 1,4,5-triphosphate), which are used in the formation of melanin (**Figure S29**). Other than this, *MELTF* (Melanotransferrin) genes also have differences. Kinesins and Dyneins are also involved in Melanogenesis and downstream processes. There are, in total, ten kinesins (*KIF14*, *KIF18A*, *KIF4B*, *KIF27*, *KIF11*, *KIF9*, *KIF13A*, *KIF21A*, *KIF7*, and *KIF26B*) and six dyneins (*DYNC2H1*, *DYNC1LI1*, *DNAAF1*, *DNAAF5*, *DRC3*, and *DYNC2LI1*), which have amino acid differences. Apart from this, *HPGDS*, which is an essential gene in pheomelanin formation, also has amino acid differences (**Table S4 and S8**).

Pathways regulating the connective tissue matrix and epidermal structures are also crucial in building and differentiating the epidermal structures. Pathways such as Extracellular matrix interaction (ECM), actin cytoskeleton, and focal adhesion play an essential role in determining and maintaining the tissue structures. ECM has amino acid differences in 11 genes, which include *AGRN* (Agrinin), *COL4A2* (Collagen), *GP1BB* (Platelet glycoprotein 1 BB), *HSPG2* (Heparan sulfate Proteoglycan 2), *ITGB1* (Integrin Beta 1), Laminins (*LAMA4*, *LAMA5*, *LAMC1*), *SDC1* (Syndecan 1), *THBS1* (Thrombospondin 1) and *VWF* (Von Willebrand Factor). In totality, there are differences in multiple laminins, collagens, glycoproteins, and other factors which are instrumental in the formation and regulation of the extracellular matrix (**Table S8 and Figure S33**). The actin cytoskeleton pathway has amino differences in 10 genes, including Actinins (*ACTN1* and *ACTN4*), *APC*, *ITGB1*, *MYLK* (Myosin light chain kinase), *NCKAP1L*, Slingshot protein phosphatase (*SSH2* and *SSH3*) and *VCL* (Vinculin) (**Table S8 and Figure S31**). The differences in these genes hint toward the differing compositions of the actin cytoskeletal system, which might interplay with ECM and focal adhesion pathway. The focal adhesion pathway has 14 amino acid difference containing genes, including multiple genes discussed above. Apart from those, *KDR*, *VEGFC*, and *ZYX* have single amino acid differences (**Table S8 and Figure S32**). The abundance of amino acid differences in the patterning pathways may reflect the role of multiple genes causing phenotypic divergence between the two species of peafowl.

**Supplementary Methods:**

**Genome annotation**

We downloaded the *Pavo muticus* genome [1], and the associated SRA reads from NCBI (**Table S1**). The genome was annotated using the Maker 3 [2] annotation pipeline. The transcriptomic data from *Pavo cristatus* was assembled using Trinity version 2.13.2 [3]. The obtained transcriptome assembly was used as EST evidence for the annotation. Protein datasets from Galliformes and UniProt curated sets were used as protein homology evidence. The first round of the Maker pipeline was executed for obtaining homology-based annotations. For the second round, de novo annotation tools such as AUGUSTUS [4] and SNAP [5] were used. Another subsequent round of annotation was performed to get the final sets of gene annotations. Gene annotation pipelines have multiple ORF with false structures and other errors [6]. We used the annotations to obtain the candidate genes and verified the ORF using the *Gallus gallus* (galgal7) genome to overcome the annotation errors.

**Gender identification**

We used Bedtools to calculate the number of reads in 50 kb windows for each bam file. To identify gender, we estimated the normalised read counts (i.e., number of reads in a 50 kb window/ mean of the number of reads in all 50 kb windows) of autosomes, Z and W chromosomes for each individual (**Supplementary Figure S1-S10**). Normalised read counts of autosomes should always be ~1, whereas, for the Z chromosome, it should be ~1 in the case of males and around ~0.5 for females. Similarly, the W chromosome should have negligible normalised read count in the case of males, whereas close to 1 in the case of females.

Based on a molecular assay, *P. cristatus* individual-1 (SAMN05660020) gender is known to be male. Keeping this individual as a reference, we divided the normalised read counts in each 50 kb window of *P. cristatus* individual-1 by the normalised read counts in the same 50 kb window in each of the other individuals. This ratio of normalised read counts of one individual's autosomes to another's autosomes will always be ~1. Since *P. cristatus* individual-1 is male, the ratio of the Z chromosome with any other individual should be 1 in the case of a male, whereas it should be around 2 in the case of a female. Using these two approaches, we identified the genders of previously unidentified individuals (see **Table S1**).

**Population genomic summary statistics:**

At present, three genome assemblies are available for *P. cristatus* [7–9] and two for *P. muticus* [1,10]; among these five genomes, the most contiguous high-quality genome is used for all the analyses. We mapped sequencing reads of 6 samples each from *P. cristatus* and *P. muticus* to the genome sequence of *P. Muticus* [10]. Genomic and transcriptomic reads were mapped using bwa mem [11] and tophat2 [12], respectively. The mapped reads were sorted and indexed using samtools [13]. We use these alignments for identifying Sample allele frequencies (SAF) which were computed using genotype likelihoods generated by ANGSD v 0.935-53-gf475f10 [14] for both peacock species separately using respective alignments. The SAF output obtained the site frequency spectrum (SFS) for both species. Sites that were covered by at least ten reads in each of the six individuals in both species were used to generate the two-dimensional SFS using the realSFS module of ANGSD [14]. The 2D-SFS between *P. cristatus* and *P. muticus* was used to generate per base, 1KB and 50 kb non-sliding window $F_{ST}$ using the FST stats2 module of ANGSD [14]. We calculated the mean estimates of population genomic statistics such as Watterson's theta ($\theta_w$), Pairwise theta ($\theta_\pi$), Tajima's D ($\tau$) and $F_{ST}$ for autosomes and the Z chromosome separately. We estimated these statistics in 1 KB windows with varying levels (at least 400 to 900 sites covered) of missing data to ensure that data quality does not affect our estimates (**Supplementary Figure S11-S20**).

Initially, we used all 12 individuals to calculate the diversity of the Z/A ratio. However, the female samples (ZW) can adversely affect the estimates of diversity for the Z chromosome. Hence, we used only male samples (3 individuals each of *P. muticus* and *P. cristatus*) to estimate genetic diversity and calculate the Z/A ratios. Similar to the initial estimates, the mean diversity (Watterson's theta and Pairwise theta) for each cutoff was used to get a mean estimate of diversity for autosomes and Z chromosomes separately.

**Shortlisting of the candidate genes**

Out of 41,183,127 quality passed sites used in differentiation analyses, 53,635 have $F_{ST}$ > 0.9, i.e., ~ 0.01% of sites are identified as highly differentiated between *Pavo cristatus* and *Pavo muticus* and might be fixed differences. We identified variant sites across the 12 individuals consisting of six samples from each species using Bcftools mpileup [15] followed by bcftools call command. Fixed sites identified by ANGSD were intersected with the vcf file using Bedtools [16] to get a fixed-sites vcf file. We used the fixed-sites vcf file to make an alternate fasta sequence for each species using AlternateFastaMaker of GATK v 4.2.0 [17].

We employed two approaches to identify candidate genes that can be used to evaluate our hypothesis. The first approach annotated the fixed-sites vcf file using SnpEff [18] and prioritised the candidate genes based on the gene-specific ratio of non-synonymous to synonymous changes. The fixed sites identified between the two species are distributed across exonic regions of 5,746 annotated genes. The exons were prioritised based on a) the ratio of fixed differences upon their length (i.e., fixed-site density) (101 of 200 highly differentiated genes showed >= one amino acid difference). (b) All genes on the Z chromosome (95 of 887 genes had >= one amino acid difference). (c) previously reported melanogenesis-related genes (20 of 145 genes had >= one amino acid difference) (see **Table S4**).

As part of the second approach, genes involved in KEGG signalling pathways [19] and other related genes were considered candidates. Since the phenotypic differences between the two species consisted of morphological traits mainly determined by developmental signalling pathways, we decided to use this as a filter to get close to the phenotype. The corresponding protein sequence from *Gallus gallus* (galgal7) was used for each candidate gene to query *Pavo cristatus* and *Pavo muticus* genomes using Exonerate [20] protein2genome tool. The obtained coding sequence was then translated, and the peptide sequence differences were tabulated between both peacocks (see **Table S5**).

**Demographic history**

The whole genomic short sequencing reads from both the species were aligned to the recent *Pavo muticus* genome [10]. The genomic bam alignments for each species were used to get consensus calls using samtools [13] and bcftools [15]. The consensus calls were converted to fastq format using vcfutils.pl vcf2fq with the quality filter of 25 and a coverage filter of greater than twice and less than one-third of the mean coverage. Obtained fastq calls were converted to the psmcfa file using the fq2psmcfa command of the psmc suite [21]. The resultant psmcfa file was then used to execute psmc with options -N30 -r5 -t5 -p 4+30*2+4+6+10. The psmc file was checked for a sufficient number of recombination events. The estimated trajectory of effective population size ($N_e$) was plotted with a mutation rate (u) of 1.33e-09 per site per year [22] with a generation time of four years [23] using psmc_plot.pl script of psmc.

**Divergence time estimate**

We employed two independent methods to obtain divergence time estimates. Genomic UCE-based alignments and pseudo-diploid analysis of psmc were performed to get estimates of divergence between these two species. For the UCE-based method, we first downloaded genome assemblies of *Pavo muticus, Pavo cristatus, Gallus gallus, Coturnix japonica, Bambusicola thoracicus, and Colinus virginianus*. We then followed the Phyluce-1.7.1 [24] pipeline to get concatenated alignments of 75% of representative gene trees. The bootstrapped species tree was obtained using this concatenated alignment using raxml-ng [25] after finding the appropriate evolution model using modeltest-ng [26]. The split time between Odontophoridae (*Galinus*) and Phasinidae (*Gallus*) was obtained from TimeTree [27] (46 MYA) and was used as fossil evidence to mark the tree. Baseml from PAMLv4.9f [28] was used to estimate the substitution rate for the nucleotide dataset and was used as a prior for mean substitution rate analysis. After running MCMCTree using usedata=3, the Hessian matrix was used as input for the next MCMCTree run. These estimates were used to determine rgene_gamma and sigma2_gamma values to execute the runs. We used rgene_gamma 11.1 and shape parameter of sigma2_gamma of 2.4 and finalised the divergence time estimation.

The WGS reads from both species were mapped to the *Pavo muticus* genome. The resultant fastq consensus calls obtained for psmc were converted to fasta using the seqtk [29] fq2fa module for pseudo-diploid analysis [30]. These fasta sequences from both species were merged using the seqtk [29] mergefa module. The combined fasta sequence was then converted back to fastq format and used for making a .psmcfa file using the fq2psmcfa module. Different -p parameters were chosen to get sufficient recombinations and the resolution in a particular time point. The parameters (-N25 -t5 -r5 -p "20+4*5") were finalized for psmc. The obtained effective population size ($N_e$) was plotted, and the time at which the line goes to infinity was considered the divergence estimate for those two species.

**Acoustic Analyses**

The representative calls for *Pavo cristatus* and *Pavo muticus* were retrieved from the animal call database Xeno-Canto [31]. In the case of multiple choices of recordings (i.e., for *Pavo cristatus*), calls from places to represent the whole distribution of the species and the recording quality were considered. Each call recording was imported in Raven pro 1.6.1 [32] for the analyses. From each recording, best and clean calls were considered for the measurements like delta time for waveform and the maximum frequency for spectrogram. A total of 43 calls (*Pavo cristatus* - 23 and *Pavo muticus* - 20) from 10 recordings (5 from each species) were sampled for the final representation. The measurement data were analysed in R [33]. The spectrogram view was exported with FFT window - 512, contrast - 75, and brightness - 75. The maximum energy bands from the spectrogram were identified as Dominant frequencies.

| **Table S1: SRA reads used in the study** | | | | | | |
|---|---|---|---|---|---|---|
| **Sr. No.** | **Species** | **Sample Accession** | **SRA Accession** | **Library Type** | **Gender (in SRA)** | **Remark** |
| 1 | *Pavo muticus* | SAMN17255116 | SRR13424288 | Genomic | Female | Gender based on coverage: Female |
| 2 | *Pavo muticus* | SAMN17255114 | SRR13424290 | Genomic | Female | Gender based on coverage: Female |
| 3 | *Pavo muticus* | SAMN15488465 | SRR12223809 | Genomic | ND | Gender based on coverage: Male |
| 4 | *Pavo muticus* | SAMN15488464 | SRR12223810 | Genomic | ND | Gender based on coverage: Male |
| 5 | *Pavo muticus* | SAMN15488463 | SRR12223811 | Genomic | ND | Gender based on coverage: Male |
| 6 | *Pavo muticus* | SAMN15488455 | SRR12223821 | Genomic | ND | Gender based on coverage: Male |
| 7 | *Pavo cristatus* | SAMN05660020 | SRR4068854 | Genomic | Male | Gender based on coverage: Male |
| 8 | *Pavo cristatus* | SAMN07739105 | | Genomic | Male | Used only to find fixed differences, Male |
| 9 | *Pavo cristatus* | SAMN03322586 | SRR1797848 | Transcriptomic | Female | Gender based on coverage: ND |
| 10 | *Pavo cristatus* | SAMN03322585 | SRR1797865 | Transcriptomic | Female | Gender based on coverage: ND |
| 11 | *Pavo cristatus* | SAMN03322587 | SRR1797860 | Transcriptomic | Male | Gender based on coverage: ND |
| 12 | *Pavo cristatus* | SAMN03322588 | SRR1797873 | Transcriptomic | Male | Gender based on coverage: ND |
| *** ND = Not Determined** | | | | | | |

| **Table S2: Summary statistics of acoustic measurements.** | | | |
|---|---|---|---|
| **Pavo cristatus (N=23)** | **Mean** | **SD** | **Range (Min-Max)** |
| **Call Period (sec)** | 0.653 | 0.072 | 0.520-0.769 |
| **Fundamental Frequency (Hz)** | 756.394 | 66.393 | 602.93-843.75 |
| **Dominant Frequency (Hz)** | 1488.663 | 122.945 | 1205.859-1687.5 |
| **Pavo muticus (N=20)** | | | |
| **Call Period (sec)** | 0.908 | 0.195 | 0.512-1.251 |
| **Fundamental Frequency (Hz)** | 551.25 | 70.696 | 516.797-689.062 |
| **Dominant Frequency (Hz)** | 1111.114 | 111.431 | 947.461-1378.125 |

| Table S3: Acoustic measurements used in the study. | | | | | | | |
|---|---|---|---|---|---|---|---|
| **Sr. No.** | **ID** | **Call period** | **Fundamental frequency** | **Dominant Frequency** | **Species** | **Location** | **Cat. nr.** |
| 1 | PCMH | 0.7690 | 843.75 | 1593.75 | *P. cristatus* | Maharashtra | XC124017 |
| 2 | PCMH | 0.6941 | 843.75 | 1593.75 | *P. cristatus* | Maharashtra | XC124017 |
| 3 | PCMH | 0.7000 | 843.75 | 1593.75 | *P. cristatus* | Maharashtra | XC124017 |
| 4 | PCMH | 0.7560 | 843.75 | 1687.5 | *P. cristatus* | Maharashtra | XC124017 |
| 5 | PCMH | 0.7552 | 843.75 | 1500 | *P. cristatus* | Maharashtra | XC124017 |
| 6 | PCMY | 0.6281 | 775.195 | 1550.391 | *P. cristatus* | Mysore | XC369407 |
| 7 | PCMY | 0.5319 | 775.195 | 1550.391 | *P. cristatus* | Mysore | XC369407 |
| 8 | PCMY | 0.5458 | 775.195 | 1636.523 | *P. cristatus* | Mysore | XC369407 |
| 9 | PCMY | 0.5926 | 775.195 | 1636.523 | *P. cristatus* | Mysore | XC369407 |
| 10 | PCMY | 0.6547 | 775.195 | 1636.523 | *P. cristatus* | Mysore | XC369407 |
| 11 | PCRJ | 0.6546 | 775.195 | 1464.258 | *P. cristatus* | Rajasthan | XC165857 |
| 12 | PCRJ | 0.6369 | 775.195 | 1550.391 | *P. cristatus* | Rajasthan | XC165857 |
| 13 | PCRJ | 0.6748 | 775.195 | 1464.258 | *P. cristatus* | Rajasthan | XC165857 |
| 14 | PCRJ | 0.7012 | 775.195 | 1464.258 | *P. cristatus* | Rajasthan | XC165857 |
| 15 | PCTN | 0.5614 | 689.062 | 1378.125 | *P. cristatus* | Tamilnadu | XC406117 |
| 16 | PCTN | 0.6615 | 689.062 | 1378.125 | *P. cristatus* | Tamilnadu | XC406117 |
| 17 | PCTN | 0.6655 | 689.062 | 1378.125 | *P. cristatus* | Tamilnadu | XC406117 |
| 18 | PCTN | 0.7076 | 689.062 | 1378.125 | *P. cristatus* | Tamilnadu | XC406117 |
| 19 | PCTN | 0.6628 | 689.062 | 1464.258 | *P. cristatus* | Tamilnadu | XC406117 |
| 20 | PCUK | 0.5633 | 775.195 | 1464.258 | *P. cristatus* | Uttarakhand | XC507742 |
| 21 | PCUK | 0.6575 | 689.062 | 1291.992 | *P. cristatus* | Uttarakhand | XC507742 |
| 22 | PCUK | 0.7207 | 602.93 | 1205.859 | *P. cristatus* | Uttarakhand | XC507742 |
| 23 | PCUK | 0.5197 | 689.062 | 1378.125 | *P. cristatus* | Uttarakhand | XC507742 |
| 24 | PMCB1 | 0.5956 | 516.797 | 1205.859 | *P. muticus* | Cambodia | XC100440 |
| 25 | PMCB1 | 0.5121 | 516.797 | 1119.727 | *P. muticus* | Cambodia | XC100440 |
| 26 | PMCB1 | 0.6964 | 516.797 | 1033.594 | *P. muticus* | Cambodia | XC100440 |
| 27 | PMCB1 | 0.8908 | 516.797 | 1119.727 | *P. muticus* | Cambodia | XC100440 |
| 28 | PMCB2 | 0.8722 | 516.797 | 1119.727 | *P. muticus* | Cambodia | XC100439 |
| 29 | PMCB2 | 0.8431 | 516.797 | 1033.594 | *P. muticus* | Cambodia | XC100439 |
| 30 | PMCB2 | 0.7921 | 516.797 | 947.461 | *P. muticus* | Cambodia | XC100439 |
| 31 | PMCB2 | 0.8552 | 516.797 | 1033.594 | *P. muticus* | Cambodia | XC100439 |
| 32 | PMIN | 0.8600 | 516.797 | 1119.727 | *P. muticus* | Indonesia | XC336671 |
| 33 | PMIN | 0.9058 | 516.797 | 1033.594 | *P. muticus* | Indonesia | XC336671 |
| 34 | PMIN | 0.8009 | 516.797 | 1119.727 | *P. muticus* | Indonesia | XC336671 |
| 35 | PMTH1 | 0.8276 | 516.797 | 1033.594 | *P. muticus* | Thailand | XC206560 |
| 36 | PMTH1 | 0.9271 | 516.797 | 1033.594 | *P. muticus* | Thailand | XC206560 |
| 37 | PMTH3 | 1.0518 | 516.797 | 1119.727 | *P. muticus* | Thailand | XC206558 |
| 38 | PMTH3 | 1.0212 | 516.797 | 1033.594 | *P. muticus* | Thailand | XC206558 |
| 39 | PMTH3 | 0.9230 | 516.797 | 1033.594 | *P. muticus* | Thailand | XC206558 |
| 40 | PMVI | 1.1252 | 689.062 | 1205.859 | *P. muticus* | Vietnam | XC19266 |
| 41 | PMVI | 1.2508 | 689.062 | 1119.727 | *P. muticus* | Vietnam | XC19266 |
| 42 | PMVI | 1.2275 | 689.062 | 1378.125 | *P. muticus* | Vietnam | XC19266 |
| 43 | PMVI | 1.1857 | 689.062 | 1378.125 | *P. muticus* | Vietnam | XC19266 |

| Table S4: The genes showing amino acid differences between *Pavo cristatus* and *Pavo muticus*. | | | | |
|---|---|---|---|---|
| **Sr. No.** | **Gene** | **amino acid #** | ***P. muticus*** | ***P. cristatus*** |
| **1** | AASDH | 26 | N | S |
| | AASDH | 279 | V | A |
| | AASDH | 661 | S | N |
| **2** | ABCC10 | 13 | P | L |
| | ABCC10 | 372 | L | F |
| | ABCC10 | 948 | I | T |
| | ABCC10 | 1004 | P | L |
| **3** | AKAP9 | 878 | V | M |
| | AKAP9 | 1156 | E | A |
| | AKAP9 | 1733 | G | R |
| | AKAP9 | 2711 | I | V |
| | AKAP9 | 3093 | T | S |
| **4** | ALMS1 | 214 | R | G |
| | ALMS1 | 1168 | V | L |
| | ALMS1 | 2290 | I | R |
| **5** | AMOTL1 | 831 | T | I |
| **6** | ANAPC1 | 1900 | V | M |
| **7** | ANK2 | 2422 | Y | F |
| | ANK2 | 2591 | I | M |
| | ANK2 | 2631 | G | S |
| **8** | APC | 289 | G | A |
| | APC | 954 | M | T |
| | APC | 1261 | P | Q |
| | APC | 1354 | D | H |
| **9** | ARID1B | 798 | T | A |
| **10** | BBS12 | 81 | T | I |
| | BBS12 | 431 | T | A |
| | BBS12 | 464 | K | R |
| | BBS12 | 557 | E | A |
| | BBS12 | 656 | D | G |
| **11** | BICC1 | 665 | A | T |
| **12** | BMP2K | 790 | S | G |
| | BMP2K | 799 | L | I |
| | BMP2K | 1043 | I | M |
| **13** | BOD1L1 | 1285 | S | N |
| | BOD1L1 | 1402 | E | K |
| | BOD1L1 | 1702 | G | E |
| | BOD1L1 | 2324 | T | A |
| **14** | BRAT1 | 574 | M | V |

| | | | | |
|---|---|---|---|---|
| | BRAT1 | 585 | I | V |
| | BRAT1 | 645 | T | A |
| | BRAT1 | 746 | G | D |
| | BRAT1 | 748 | H | Q |
| **15** | BRCA1 | 317 | Q | R |
| | BRCA1 | 319 | S | G |
| | BRCA1 | 812 | I | M |
| | BRCA1 | 1571 | I | V |
| **16** | BRCA2 | 694 | N | S |
| | BRCA2 | 1081 | S | N |
| | BRCA2 | 1870 | I | S |
| | BRCA2 | 1974 | H | Q |
| | BRCA2 | 2834 | V | I |
| | BRCA2 | 3305 | L | P |
| **17** | C1orf21 | 75 | S | P |
| **18** | C1R | 161 | Q | R |
| **19** | C8orf48 | 21 | W | R |
| | C8orf48 | 48 | R | H |
| | C8orf48 | 198 | R | Q |
| **20** | CCP110 | 608 | R | Q |
| | CCP110 | 984 | S | R |
| **21** | CDK5RAP2 | 338 | A | V |
| | CDK5RAP2 | 586 | V | I |
| | CDK5RAP2 | 2005 | G | S |
| | CDK5RAP2 | 2059 | N | D |
| **22** | CENPE | 782 | T | S |
| | CENPE | 1598 | D | N |
| | CENPE | 1851 | T | A |
| | CENPE | 1951 | N | T |
| | CENPE | 2015 | F | Y |
| | CENPE | 2084 | M | I |
| **23** | CENPF | 1267 | H | L |
| **24** | CEP164 | 1550 | D | E |
| **25** | CGGBP1 | 79 | I | V |
| **26** | CKAP2L | 383 | M | T |
| **27** | CMYA5 | 677 | S | C |
| | CMYA5 | 3014 | E | Q |
| **28** | COBLL1 | 361 | Y | H |
| | COBLL1 | 424 | R | C |
| **29** | COL15A1 | 282 | I | T |
| **30** | CPED1 | 376 | V | I |

| | | | | |
|---|---|---|---|---|
| **31** | CRYBG3 | 405 | Q | H |
| | CRYBG3 | 597 | H | D |
| | CRYBG3 | 898 | I | T |
| | CRYBG3 | 907 | M | K |
| | CRYBG3 | 1427 | K | E |
| | CRYBG3 | 1505 | S | N |
| **32** | CSPG4B | 2491 | R | S |
| **33** | CUL9 | 925 | M | L |
| | CUL9 | 1968 | N | S |
| | CUL9 | 1970 | P | S |
| | CUL9 | 2010 | T | A |
| **34** | DACH2 | 290 | A | T |
| **35** | DYNC2H1 | 1079 | Q | R |
| | DYNC2H1 | 3505 | T | I |
| **36** | EP400 | 1815 | S | T |
| **37** | FAM214B | 14 | H | R |
| | FAM214B | 390 | S | A |
| | FAM217B | 483 | T | A |
| **38** | GCC1 | 128 | G | A |
| **39** | GCC2 | 291 | M | V |
| | GCC2 | 781 | D | N |
| | GCC2 | 1437 | S | G |
| | GCC2 | 1492 | A | T |
| | GCC2 | 1497 | V | L |
| | GCC2 | 1500 | S | N |
| **40** | GLTSCR1L | 960 | I | T |
| **41** | GSAP | 292 | A | T |
| **42** | HABP4 | 98 | E | Q |
| | HABP4 | 100 | D | E |
| | HABP4 | 155 | R | H |
| **43** | HELQ | 577 | K | E |
| **44** | HMCN1 | 1851 | V | I |
| | HMCN1 | 2030 | I | V |
| | HMCN1 | 2244 | S | L |
| | HMCN1 | 2307 | A | T |
| | HMCN1 | 3456 | G | S |
| | HMCN1 | 4582 | R | C |
| | HMCN1 | 4948 | N | S |
| | HMCN1 | 5315 | S | A |
| **45** | IGF2R | 753 | V | I |
| | IGF2R | 920 | R | Q |

| | | | | |
|---|---|---|---|---|
| | IGF2R | 1878 | L | F |
| **46** | KAT6B | 1348 | E | Q |
| **47** | KCTD20 | 57 | N | S |
| **48** | KDELC2 | 80 | I | T |
| **49** | KIF7 | 695 | L | S |
| | KIF7 | 1293 | T | A |
| **50** | KMT2A | 2242 | V | G |
| | KMT2A | 2722 | A | V |
| | KMT2A | 3306 | A | T |
| **51** | LACTB2 | 276 | C | R |
| **52** | LOC10175063[illegible] | 797 | V | A |
| **53** | LTBP1 | 309 | G | S |
| **54** | LTK | 436 | T | I |
| **55** | MBTPS1 | 124 | M | T |
| | MBTPS1 | 672 | M | V |
| **56** | METTL2B | 133 | M | T |
| **57** | MKI67 | 190 | A | E |
| | MKI67 | 428 | V | A |
| | MKI67 | 847 | N | S |
| | MKI67 | 1045 | G | D |
| | MKI67 | 1542 | G | E |
| | MKI67 | 1868 | N | S |
| | MKI67 | 1934 | I | M |
| **58** | MXRA5 | 897 | V | L |
| | MXRA5 | 1435 | P | R |
| | MXRA5 | 1495 | A | V |
| | MXRA5 | 2560 | Q | R |
| **59** | NBAS | 286 | Y | S |
| | NBAS | 1135 | V | I |
| **60** | NBEAL1 | 1451 | N | D |
| | NBEAL1 | 2224 | Y | H |
| **61** | NBR1 | 327 | I | V |
| | NRP2 | 846 | L | S |
| **62** | PALB2 | 95 | R | G |
| **63** | PCNT | 379 | V | I |
| | PCNT | 460 | N | D |
| | PCNT | 931 | V | M |
| | PCNT | 1054 | I | V |
| | PCNT | 1935 | F | L |
| | PCNT | 2028 | K | R |
| | PCNT | 2584 | R | C |

| | | | | |
|---|---|---|---|---|
| **64** | PDZD2 | 1523 | T | M |
| | PDZD2 | 1662 | T | A |
| | PDZD2 | 1735 | V | L |
| **65** | PHYKPL | 372 | I | V |
| **66** | RALB | 51 | R | S |
| **67** | RNF213 | 1199 | I | V |
| | RNF213 | 3024 | V | I |
| | RNF213 | 4420 | S | G |
| **68** | RNF217 | 353 | S | T |
| **69** | RTEL1 | 921 | T | A |
| **70** | RUFY3 | 536 | R | W |
| | RUFY3 | 569 | D | G |
| **71** | SECISBP2 | 227 | L | S |
| | SECISBP2 | 271 | S | L |
| | SECISBP2 | 440 | F | I |
| | SECISBP2 | 455 | R | W |
| | SECISBP2 | 881 | K | E |
| **72** | SH3TC2 | 258 | E | V |
| **73** | SIK1 | 511 | S | T |
| **74** | SIK2 | 298 | I | V |
| | SIK2 | 528 | V | M |
| **75** | SIMC1 | 692 | V | A |
| **76** | SLC44A5 | 586 | D | E |
| **77** | SLF2 | 428 | D | N |
| | SLF2 | 434 | E | K |
| | SLF2 | 442 | A | D |
| | SLF2 | 452 | S | N |
| | SLF2 | 901 | Q | R |
| **78** | SMS | 75 | S | N |
| | SMS | 77 | S | N |
| **79** | SON | 851 | I | T |
| | SON | 1186 | M | V |
| | SON | 1302 | K | M |
| | SON | 1355 | T | A |
| | SON | 1469 | I | V |
| **80** | SPICE1 | 304 | A | T |
| | SPICE1 | 376 | C | S |
| | SPICE1 | 752 | V | A |
| | SPICE1 | 788 | R | T |
| | SPICE1 | 859 | T | I |
| **81** | STON1 | 44 | H | P |

| | | | | |
|---|---|---|---|---|
| | STON1 | 191 | N | S |
| | STON1 | 267 | W | R |
| **82** | TDRD3 | 561 | Q | H |
| | TDRD3 | 589 | H | N |
| **83** | TET2 | 149 | C | R |
| | TET2 | 234 | A | T |
| | TET2 | 605 | H | Q |
| | TET2 | 791 | Q | H |
| **84** | TGS1 | 178 | V | I |
| | TGS1 | 284 | M | V |
| | TGS1 | 298 | L | P |
| | TGS1 | 300 | T | A |
| | TGS1 | 310 | P | L |
| | TGS1 | 329 | D | H |
| | TGS1 | 335 | V | I |
| | TGS1 | 607 | T | I |
| **85** | TJP3 | 387 | E | D |
| | TJP3 | 547 | L | M |
| | TJP3 | 556 | H | R |
| **86** | TMEM14C | 68 | V | I |
| **87** | TMEM38B | 73 | E | A |
| **88** | TRAF6 | 45 | L | P |
| **89** | UACA | 282 | E | D |
| | UACA | 658 | S | F |
| **90** | UIMC1 | 667 | V | D |
| **91** | USP47 | 175 | A | S |
| **92** | USPL1 | 521 | R | Q |
| **93** | VCPIP1 | 760 | T | A |
| **94** | VPS13B | 2051 | F | I |
| | VPS13B | 2386 | S | T |
| **95** | YAE1D1 | 173 | G | S |
| | YAE1D1 | 178 | D | G |
| | YAE1D1 | 181 | G | R |
| **96** | ZBTB21 | 287 | R | K |
| | ZBTB21 | 429 | T | S |
| **97** | ZNF318 | 172 | L | P |
| | ZNF318 | 672 | L | P |
| | ZNF318 | 689 | I | V |
| | ZNF318 | 1863 | S | P |
| **98** | ZNF407 | 223 | K | R |
| | ZNF407 | 498 | H | L |

| | | | | |
|---|---|---|---|---|
| | ZNF407 | 711 | I | V |
| | ZNF407 | 991 | C | S |
| | ZNF407 | 1103 | T | A |
| **99** | ZNF410 | 163 | A | T |
| **100** | ZNF462 | 410 | T | A |
| **101** | ZNF648 | 253 | A | T |
| **Melanogenesis related genes** | | | | |
| **Sr. No.** | **Gene** | **amino acid #** | ***P. muticus*** | ***P. cristatus*** |
| **1** | LOC416959 | 478 | R | H |
| | LOC416959 | 735 | V | I |
| **2** | MELTF | 179 | S | N |
| **3** | KIF14 | 71 | C | F |
| | KIF14 | 1409 | H | R |
| **4** | KIF18A | 524 | R | H |
| **5** | KIF4B | 616 | N | S |
| | KIF4B | 751 | N | S |
| **6** | KIF27 | 566 | D | N |
| **7** | KIF11 | 846 | Q | R |
| | KIF11 | 1188 | E | D |
| **8** | KIF9 | 393 | V | L |
| | KIF9 | 502 | F | L |
| **9** | KIF13A | 1612 | G | S |
| **10** | KIF21A | 585 | R | K |
| | KIF21A | 1608 | M | V |
| **11** | KIF7 | 695 | L | S |
| | KIF7 | 1293 | T | A |
| **12** | KIF26B | 583 | V | L |
| **13** | DYNC2H1 | 1079 | Q | R |
| | DYNC2H1 | 3505 | T | I |
| **14** | DYNC1LI1 | 157 | N | S |
| **15** | DNAAF1 | 55 | C | G |
| **16** | DNAAF5 | 744 | A | T |
| | DNAAF5 | 827 | Q | R |
| **17** | DRC3 | 515 | T | I |
| **18** | DYNC2LI1 | 178 | T | I |
| **19** | HPGDS | 37 | V | A |
| **20** | PLCB1 | 908 | L | S |
| **21** | CREB3L4 | 339 | A | T |
| **Chr Z genes** | | | | |

| | | | | |
|---|---|---|---|---|
| **1** | ADAMTS12 | 1257 | V | L |
| **2** | AGGF1 | 111 | V | A |
| **3** | ANKRD31 | 394 | C | S |
| | ANKRD31 | 1354 | N | S |
| | ANKRD31 | 1449 | H | R |
| **4** | AOPEP | 370 | L | P |
| **5** | APC | 289 | G | A |
| | APC | 954 | M | T |
| | APC | 1261 | P | Q |
| | APC | 1354 | D | H |
| **6** | ARSB | 450 | M | T |
| **7** | BDP1 | 871 | S | A |
| | BDP1 | 1389 | S | P |
| | BDP1 | 2029 | D | E |
| | BDP1 | 2098 | T | I |
| **8** | C6 | 192 | R | C |
| | C6 | 231 | L | S |
| | C6 | 711 | H | R |
| **9** | CCDC112 | 263 | E | K |
| **10** | CCDC125 | 116 | T | I |
| **11** | CCDC171 | 919 | N | S |
| | CCDC171 | 1157 | S | R |
| **12** | CEP120 | 450 | A | V |
| | CEP120 | 830 | E | Q |
| **13** | CEP78 | 373 | L | V |
| | CEP78 | 382 | I | L |
| **14** | CMYA5 | 677 | S | C |
| | CMYA5 | 3014 | E | Q |
| **15** | CNTLN | 256 | T | A |
| | CNTLN | 725 | R | C |
| | CNTLN | 943 | T | I |
| **16** | CPLANE1 | 2982 | M | V |
| **17** | CR1 | 1793 | L | P |
| | CR1 | 2509 | Y | H |
| | CR1 | 2869 | V | I |
| | CR1 | 3223 | V | I |
| **18** | CREB3L4 | 339 | A | T |
| **19** | CSPG4B | 2491 | R | S |
| **20** | DGKQ | 679 | I | V |
| **21** | DIMT1 | 118 | V | A |
| **22** | DOCK8 | 1959 | Q | R |

| | | | | |
|---|---|---|---|---|
| **23** | EPG5 | 2021 | G | D |
| **24** | ERCC6L2 | 192 | R | C |
| | ERCC6L2 | 231 | L | S |
| | ERCC6L2 | 711 | H | R |
| **25** | FAM214B | 14 | H | R |
| | FAM214B | 390 | S | A |
| **26** | FBN2 | 419 | D | G |
| **27** | FYB1 | 435 | I | V |
| | FYB1 | 739 | P | S |
| **28** | GAK | 803 | H | R |
| | GAK | 1051 | G | D |
| **29** | GFM2 | 160 | T | A |
| **30** | GRAMD3 | 405 | T | I |
| **31** | GRHPR | 318 | E | K |
| **32** | HABP4 | 98 | E | Q |
| | HABP4 | 100 | D | E |
| | HABP4 | 155 | R | H |
| **33** | HAUS1 | 47 | F | I |
| **34** | HAUS6 | 98 | E | K |
| | HAUS6 | 258 | H | R |
| | HAUS6 | 361 | E | K |
| **35** | IDNK | 129 | I | V |
| **36** | IKBKAP | 648 | S | N |
| **37** | IL6ST | 14 | A | V |
| | IL6ST | 379 | S | T |
| **38** | KANK1 | 1052 | P | A |
| **39** | KIAA0368 | 209 | S | T |
| **40** | KIAA2026 | 1279 | T | A |
| | KIAA2026 | 1879 | T | A |
| | KIAA2026 | 1961 | V | M |
| **41** | KIF27 | 566 | D | N |
| **42** | LIFR | 101 | A | T |
| **43** | LNPEP | 567 | M | T |
| | LNPEP | 1013 | T | I |
| **44** | LOC112530730 | 48 | R | Q |
| | LOC112530730 | 60 | N | D |
| **41** | LOC112530732 | 48 | R | Q |
| | LOC112530732 | 60 | N | D |
| **42** | LOC112530763 | 60 | R | Q |
| | LOC112530763 | 63 | G | R |
| **43** | LOC407092 | 158 | N | S |

| | | | | |
|---|---|---|---|---|
| **44** | MAP1B | 1038 | Y | C |
| **45** | MOCS2 | 66 | I | M |
| **46** | MPDZ | 1043 | R | P |
| | MPDZ | 1070 | S | P |
| | MPDZ | 1937 | A | G |
| **47** | MTMR12 | 656 | I | M |
| | MTMR12 | 686 | N | S |
| **48** | NANS | 141 | P | L |
| **49** | NOL6 | 308 | N | S |
| **50** | NUDT12 | 181 | L | H |
| | NUDT12 | 189 | S | N |
| **51** | PALM2AKAP2 | 591 | N | D |
| | PALM2AKAP2 | 724 | S | N |
| **52** | PDZD2 | 1523 | T | M |
| | PDZD2 | 1662 | T | A |
| | PDZD2 | 1735 | V | L |
| **53** | PIGG | 463 | S | C |
| | PIGG | 621 | S | R |
| **54** | PLAA | 767 | F | V |
| **55** | POC5 | 240 | H | R |
| **56** | POLK | 15 | S | * |
| **57** | PTAR1 | 362 | R | H |
| | PTAR1 | 434 | D | G |
| **58** | PUM3 | 137 | M | V |
| | PUM3 | 600 | R | Q |
| **59** | RIOK2 | 404 | L | V |
| **60** | RLN3 | 160 | S | T |
| **67** | RMI1 | 396 | R | Q |
| | RMI1 | 441 | I | S |
| **68** | ROR2 | 149 | H | Y |
| **69** | RUSC2 | 565 | T | A |
| | RUSC2 | 807 | S | N |
| **70** | SECISBP2 | 227 | L | S |
| | SECISBP2 | 271 | S | L |
| | SECISBP2 | 440 | F | I |
| | SECISBP2 | 455 | R | W |
| | SECISBP2 | 881 | K | E |
| **71** | SHLD3 | 226 | I | L |
| **72** | SLC24A2 | 23 | G | D |
| **73** | SLC25A51 | 93 | G | R |
| **74** | SLC44A1 | 264 | Q | R |

| | | | | |
|---|---|---|---|---|
| **75** | SLF1 | 643 | M | V |
| **76** | SLF2 | 428 | D | N |
| | SLF2 | 434 | E | K |
| | SLF2 | 442 | A | D |
| | SLF2 | 452 | S | N |
| | SLF2 | 901 | Q | R |
| **77** | SNCAIP | 849 | M | L |
| | SNCAIP | 939 | V | I |
| **78** | SUSD1 | 233 | V | I |
| | SUSD1 | 278 | Q | H |
| **79** | SYK | 10 | S | N |
| **80** | TAF1C | 263 | R | C |
| **81** | TBC1D2 | 96 | S | Y |
| **82** | TMC1 | 17 | A | P |
| | TMC1 | 912 | P | L |
| **83** | TMEM38B | 73 | E | A |
| **84** | TMEM8B | 463 | H | R |
| **85** | TRIM36 | 559 | S | T |
| | TRIM36 | 662 | S | A |
| **86** | TTC33 | 197 | E | Q |
| **87** | UBQLN1 | 125 | S | T |
| **88** | VPS13A | 2044 | S | G |
| | VPS13A | 2404 | E | G |
| **89** | WDR36 | 733 | T | A |
| **90** | WDR41 | 419 | A | V |
| **91** | ZBTB5 | 175 | I | M |
| **92** | ZCCHC6 | 668 | C | Y |
| | ZCCHC6 | 673 | T | K |
| **93** | ZDHHC21 | 48 | V | L |
| **94** | ZFYVE16 | 31 | A | T |
| | ZFYVE16 | 282 | C | Y |
| | ZFYVE16 | 335 | S | F |
| | ZFYVE16 | 599 | Q | E |
| **95** | ZNF462 | 410 | T | A |

**Table S5: Mean values and Z/A ratios of Watterson's theta (Tw), Pairwise theta (Tp), Tajima's D and FST for autosomes and Z chromosome of *P. cristatus* and *P. muticus* in 1KB windows.**

**Watterson's Theta**

| | | | | | *P. muticus* | | | | | |
|---|---|---|---|---|---|---|---|---|---|---|
| | | AUTOSOMES | | | | | | Z chromosome | | |
| Sr. No. | Nsites | Number of windows | Total sites (in MB) | Tw (mean) | Z/A ratio | Sr. No. | Nsites | Number of windows | Total sites (in MB) | Tw (mean) |
| 1 | 400 | 838347 | 838.347 | 0.00128 | 0.54326 | 1 | 400 | 56064 | 56.064 | 0.00070 |
| 2 | 500 | 765727 | 765.727 | 0.00126 | 0.55003 | 2 | 500 | 45068 | 45.068 | 0.00070 |
| 3 | 600 | 650050 | 650.05 | 0.00125 | 0.55747 | 3 | 600 | 32086 | 32.086 | 0.00070 |
| 4 | 700 | 488797 | 488.797 | 0.00123 | 0.56340 | 4 | 700 | 19078 | 19.078 | 0.00069 |
| 5 | 800 | 301829 | 301.829 | 0.00121 | 0.58128 | 5 | 800 | 8662 | 8.662 | 0.00071 |
| 6 | 900 | 129149 | 129.149 | 0.00120 | 0.61006 | 6 | 900 | 2392 | 2.392 | 0.00073 |
| | | | Mean | 0.00124 | 0.56758 | | | | Mean | 0.00070 |

| | | | | | *P. cristatus* | | | | | |
|---|---|---|---|---|---|---|---|---|---|---|
| | | AUTOSOMES | | | | | | Z chromosome | | |
| Sr. No. | Nsites | Number of windows | Total sites (in MB) | Tw (mean) | Z/A ratio | Sr. No. | Nsites | Number of windows | Total sites (in MB) | Tw (mean) |
| 1 | 400 | 36362 | 36.362 | 0.00152 | 0.44559 | 1 | 400 | 1410 | 1.41 | 0.00068 |
| 2 | 500 | 28595 | 28.595 | 0.00151 | 0.43561 | 2 | 500 | 1117 | 1.117 | 0.00066 |
| 3 | 600 | 22376 | 22.376 | 0.00151 | 0.42975 | 3 | 600 | 882 | 0.882 | 0.00065 |
| 4 | 700 | 17091 | 17.091 | 0.00150 | 0.40461 | 4 | 700 | 690 | 0.69 | 0.00061 |
| 5 | 800 | 12397 | 12.397 | 0.00147 | 0.42275 | 5 | 800 | 501 | 0.501 | 0.00062 |
| 6 | 900 | 7723 | 7.723 | 0.00145 | 0.40969 | 6 | 900 | 322 | 0.322 | 0.00060 |
| | | | Mean | 0.00149 | 0.42467 | | | | Mean | 0.00064 |

**Pairwise Theta**

| | | | | | *P. muticus* | | | | | |
|---|---|---|---|---|---|---|---|---|---|---|
| | | AUTOSOMES | | | | | | Z chromosome | | |
| Sr. No. | Nsites | Number of windows | Total sites (in MB) | Tp (mean) | Z/A ratio | Sr. No. | Nsites | Number of windows | Total sites (in MB) | Tp (mean) |
| 1 | 400 | 838347 | 838.347 | 0.00120 | 0.56796 | 1 | 400 | 56064 | 56.064 | 0.00068 |
| 2 | 500 | 765727 | 765.727 | 0.00118 | 0.57623 | 2 | 500 | 45068 | 45.068 | 0.00068 |
| 3 | 600 | 650050 | 650.05 | 0.00117 | 0.58458 | 3 | 600 | 32086 | 32.086 | 0.00068 |
| 4 | 700 | 488797 | 488.797 | 0.00115 | 0.59093 | 4 | 700 | 19078 | 19.078 | 0.00068 |
| 5 | 800 | 301829 | 301.829 | 0.00113 | 0.61127 | 5 | 800 | 8662 | 8.662 | 0.00069 |
| 6 | 900 | 129149 | 129.149 | 0.00112 | 0.63739 | 6 | 900 | 2392 | 2.392 | 0.00072 |
| | | | Mean | 0.00116 | 0.59473 | | | | Mean | 0.00069 |

| | | | | | *P. cristatus* | | | | | |
|---|---|---|---|---|---|---|---|---|---|---|
| | | AUTOSOMES | | | | | | Z chromosome | | |
| Sr. No. | Nsites | Number of windows | Total sites (in MB) | Tp (mean) | Z/A ratio | Sr. No. | Nsites | Number of windows | Total sites (in MB) | Tp (mean) |
| 1 | 400 | 36362 | 36.362 | 0.00153 | 0.41724 | 1 | 400 | 1410 | 1.41 | 0.00064 |
| 2 | 500 | 28595 | 28.595 | 0.00153 | 0.40944 | 2 | 500 | 1117 | 1.117 | 0.00063 |
| 3 | 600 | 22376 | 22.376 | 0.00152 | 0.40313 | 3 | 600 | 882 | 0.882 | 0.00061 |
| 4 | 700 | 17091 | 17.091 | 0.00152 | 0.37215 | 4 | 700 | 690 | 0.69 | 0.00057 |
| 5 | 800 | 12397 | 12.397 | 0.00150 | 0.38936 | 5 | 800 | 501 | 0.501 | 0.00058 |
| 6 | 900 | 7723 | 7.723 | 0.00148 | 0.37405 | 6 | 900 | 322 | 0.322 | 0.00055 |
| | | | Mean | 0.00151 | 0.39423 | | | | Mean | 0.00060 |

**FST**

| | | AUTOSOMES | | | | | | Z chromosome | | |
|---|---|---|---|---|---|---|---|---|---|---|
| Sr. No. | Nsites | Number of windows | Total sites (in MB) | FST (mean) | | Sr. No. | Nsites | Number of windows | Total sites (in MB) | FST (mean) |
| 1 | 400 | 22562 | 22.562 | 0.55000 | | 1 | 400 | 630 | 0.63 | 0.64000 |
| 2 | 500 | 15784 | 15.784 | 0.57000 | | 2 | 500 | 403 | 0.403 | 0.67000 |
| 3 | 600 | 10324 | 10.324 | 0.59000 | | 3 | 600 | 218 | 0.218 | 0.69000 |
| 4 | 700 | 6163 | 6.163 | 0.60000 | | 4 | 700 | 112 | 0.112 | 0.72000 |
| 5 | 800 | 2948 | 2.948 | 0.61000 | | 5 | 800 | 37 | 0.037 | 0.79000 |
| 6 | 900 | 910 | 0.91 | 0.62000 | | 6 | 900 | 11 | 0.011 | 0.85000 |
| | | | Mean | 0.59000 | | | | | Mean | 0.72667 |

**Tajima's D**

| | | | | | *P. muticus* | | | | | |
|---|---|---|---|---|---|---|---|---|---|---|
| | | AUTOSOMES | | | | | | Z chromosome | | |
| Sr. No. | Nsites | Number of windows | Total sites (in MB) | TD (mean) | | Sr. No. | Nsites | Number of windows | Total sites (in MB) | TD (mean) |
| 1 | 400 | 838350 | 838.35 | -0.21754 | | 1 | 400 | 56064 | 56.064 | -0.09762 |
| 2 | 500 | 765728 | 765.728 | -0.22139 | | 2 | 500 | 45068 | 45.068 | -0.09795 |
| 3 | 600 | 650051 | 650.051 | -0.22571 | | 3 | 600 | 32086 | 32.086 | -0.10076 |
| 4 | 700 | 488798 | 488.798 | -0.23018 | | 4 | 700 | 19078 | 19.078 | -0.10279 |

| | | | | | | | | | | |
|---|---|---|---|---|---|---|---|---|---|---|
| 5 | 800 | 301830 | 301.83 | -0.23653 | | 5 | 800 | 8662 | 8.662 | -0.10450 |
| 6 | 900 | 129150 | 129.15 | -0.24012 | -0.22883 | 6 | 900 | 2392 | 2.392 | -0.12899 |
| | | | | | ***P. cristatus*** | | | | | |
| | | **AUTOSOMES** | | | | | | **Z chromosome** | | |
| **Sr. No.** | **Nsites** | **Number of windows** | **Total sites (in MB)** | **TD (mean)** | | **Sr. No.** | **Nsites** | **Number of windows** | **Total sites (in MB)** | **TD (mean)** |
| 1 | 400 | 37186 | 37.186 | 0.01885 | | 1 | 400 | 1505 | 1.505 | -0.09266 |
| 2 | 500 | 29121 | 29.121 | 0.01913 | | 2 | 500 | 1180 | 1.18 | -0.09093 |
| 3 | 600 | 22751 | 22.751 | 0.02331 | | 3 | 600 | 928 | 0.928 | -0.10006 |
| 4 | 700 | 17390 | 17.39 | 0.03051 | | 4 | 700 | 726 | 0.726 | -0.11976 |
| 5 | 800 | 12638 | 12.638 | 0.04426 | | 5 | 800 | 531 | 0.531 | -0.10330 |
| 6 | 900 | 7920 | 7.92 | 0.05767 | 0.03826 | 6 | 900 | 344 | 0.344 | -0.12170 |

**Table S6: Male sample based estimates of mean values and Z/A ratios of Watterson's theta (Tw), Pairwise theta (Tp), Tajima's D and FST for autosomes and Z chromosome of *P. cristatus* and *P. muticus* in 1KB windows.**

**Watterson's Theta**

| | | | | | *P. muticus* | | | | | |
|---|---|---|---|---|---|---|---|---|---|---|
| | | AUTOSOMES | | | | | | Z chromosome | | |
| Sr. No. | Nsites | Number of windows | Total sites (in MB) | Tw (mean) | Z/A ratio | Sr. No. | Nsites | Number of windows | Total sites (in MB) | Tw (mean) |
| 1 | 400 | 918328 | 918.328 | 0.00127 | 0.57788 | 1 | 400 | 73457 | 73.457 | 0.00073 |
| 2 | 500 | 913492 | 913.492 | 0.00126 | 0.57888 | 2 | 500 | 73178 | 73.178 | 0.00073 |
| 3 | 600 | 906886 | 906.886 | 0.00125 | 0.58011 | 3 | 600 | 72725 | 72.725 | 0.00073 |
| 4 | 700 | 895281 | 895.281 | 0.00124 | 0.58204 | 4 | 700 | 71815 | 71.815 | 0.00072 |
| 5 | 800 | 865510 | 865.51 | 0.00123 | 0.58417 | 5 | 800 | 69424 | 69.424 | 0.00072 |
| 6 | 900 | 772708 | 772.708 | 0.00122 | 0.58856 | 6 | 900 | 61870 | 61.87 | 0.00072 |
| | | | Mean | 0.00125 | 0.58194 | | | | Mean | 0.00072 |

| | | | | | *P. cristatus* | | | | | |
|---|---|---|---|---|---|---|---|---|---|---|
| | | AUTOSOMES | | | | | | Z chromosome | | |
| Sr. No. | Nsites | Number of windows | Total sites (in MB) | Tw (mean) | Z/A ratio | Sr. No. | Nsites | Number of windows | Total sites (in MB) | Tw (mean) |
| 1 | 400 | 113658 | 113.658 | 0.00194 | 0.44806 | 1 | 400 | 6351 | 6.351 | 0.00087 |
| 2 | 500 | 93401 | 93.401 | 0.00195 | 0.44403 | 2 | 500 | 5169 | 5.169 | 0.00087 |
| 3 | 600 | 75698 | 75.698 | 0.00195 | 0.44789 | 3 | 600 | 4157 | 4.157 | 0.00087 |
| 4 | 700 | 59860 | 59.86 | 0.00194 | 0.43928 | 4 | 700 | 3216 | 3.216 | 0.00085 |
| 5 | 800 | 44833 | 44.833 | 0.00192 | 0.45184 | 5 | 800 | 2354 | 2.354 | 0.00087 |
| 6 | 900 | 29709 | 29.709 | 0.00187 | 0.45117 | 6 | 900 | 1510 | 1.51 | 0.00085 |
| | | | Mean | 0.00193 | 0.44705 | | | | Mean | 0.00086 |

**Pairwise Theta**

| | | | | | *P. muticus* | | | | | |
|---|---|---|---|---|---|---|---|---|---|---|
| | | AUTOSOMES | | | | | | Z chromosome | | |
| Sr. No. | Nsites | Number of windows | Total sites (in MB) | Tp (mean) | Z/A ratio | Sr. No. | Nsites | Number of windows | Total sites (in MB) | Tp (mean) |
| 1 | 400 | 918328 | 918.328 | 0.00125 | 0.57527 | 1 | 400 | 73457 | 73.457 | 0.00072 |
| 2 | 500 | 913492 | 913.492 | 0.00124 | 0.57625 | 2 | 500 | 73178 | 73.178 | 0.00072 |
| 3 | 600 | 906886 | 906.886 | 0.00124 | 0.57736 | 3 | 600 | 72725 | 72.725 | 0.00071 |
| 4 | 700 | 895281 | 895.281 | 0.00123 | 0.57926 | 4 | 700 | 71815 | 71.815 | 0.00071 |
| 5 | 800 | 865510 | 865.51 | 0.00121 | 0.58111 | 5 | 800 | 69424 | 69.424 | 0.00071 |
| 6 | 900 | 772708 | 772.708 | 0.00120 | 0.58544 | 6 | 900 | 61870 | 61.87 | 0.00070 |
| | | | Mean | 0.00123 | 0.57911 | | | | Mean | 0.00071 |

| | | | | | *P. cristatus* | | | | | |
|---|---|---|---|---|---|---|---|---|---|---|
| | | AUTOSOMES | | | | | | Z chromosome | | |
| Sr. No. | Nsites | Number of windows | Total sites (in MB) | Tp (mean) | Z/A ratio | Sr. No. | Nsites | Number of windows | Total sites (in MB) | Tp (mean) |
| 1 | 400 | 113658 | 113.658 | 0.00193 | 0.43390 | 1 | 400 | 6351 | 6.351 | 0.00084 |
| 2 | 500 | 93401 | 93.401 | 0.00194 | 0.42994 | 2 | 500 | 5169 | 5.169 | 0.00083 |
| 3 | 600 | 75698 | 75.698 | 0.00194 | 0.43296 | 3 | 600 | 4157 | 4.157 | 0.00084 |
| 4 | 700 | 59860 | 59.86 | 0.00193 | 0.42392 | 4 | 700 | 3216 | 3.216 | 0.00082 |
| 5 | 800 | 44833 | 44.833 | 0.00191 | 0.43551 | 5 | 800 | 2354 | 2.354 | 0.00083 |
| 6 | 900 | 29709 | 29.709 | 0.00187 | 0.43671 | 6 | 900 | 1510 | 1.51 | 0.00082 |
| | | | Mean | 0.00192 | 0.43216 | | | | Mean | 0.00083 |

**FST**

| | | AUTOSOMES | | | | | | Z chromosome | | |
|---|---|---|---|---|---|---|---|---|---|---|
| Sr. No. | Nsites | Number of windows | Total sites (in MB) | FST (mean) | | Sr. No. | Nsites | Number of windows | Total sites (in MB) | FST (mean) |
| 1 | 400 | 106649 | 106.649 | 0.56725 | | 1 | 400 | 6030 | 6.03 | 0.68983 |
| 2 | 500 | 87151 | 87.151 | 0.58220 | | 2 | 500 | 4896 | 4.896 | 0.71168 |
| 3 | 600 | 70071 | 70.071 | 0.59369 | | 3 | 600 | 3922 | 3.922 | 0.72976 |
| 4 | 700 | 54470 | 54.47 | 0.60454 | | 4 | 700 | 2964 | 2.964 | 0.74687 |
| 5 | 800 | 39599 | 39.599 | 0.61431 | | 5 | 800 | 2103 | 2.103 | 0.75924 |
| 6 | 900 | 24222 | 24.222 | 0.62426 | | 6 | 900 | 1261 | 1.261 | 0.76759 |
| | | | Mean | 0.59771 | | | | | Mean | 0.73416 |

**Tajima's D**

| | | | | | *P. muticus* | | | | | |
|---|---|---|---|---|---|---|---|---|---|---|
| | | AUTOSOMES | | | | | | Z chromosome | | |
| Sr. No. | Nsites | Number of windows | Total sites (in MB) | TD (mean) | | Sr. No. | Nsites | Number of windows | Total sites (in MB) | TD (mean) |
| 1 | 400 | 918330 | 918.33 | -0.11239 | | 1 | 400 | 73457 | 73.457 | -0.11480 |
| 2 | 500 | 913494 | 913.494 | -0.11278 | | 2 | 500 | 73178 | 73.178 | -0.11510 |
| 3 | 600 | 906888 | 906.888 | -0.11322 | | 3 | 600 | 72725 | 72.725 | -0.11537 |
| 4 | 700 | 895283 | 895.283 | -0.11373 | | 4 | 700 | 71815 | 71.815 | -0.11571 |
| 5 | 800 | 865512 | 865.512 | -0.11436 | | 5 | 800 | 69424 | 69.424 | -0.11672 |
| 6 | 900 | 772710 | 772.71 | -0.11474 | -0.11357 | 6 | 900 | 61870 | 61.87 | -0.11666 |

| | | | | | *P. cristatus* | | | | | |
|---|---|---|---|---|---|---|---|---|---|---|
| | | AUTOSOMES | | | | | | Z chromosome | | |
| Sr. No. | Nsites | Number of windows | Total sites (in MB) | TD (mean) | | Sr. No. | Nsites | Number of windows | Total sites (in MB) | TD (mean) |
| 1 | 400 | 114701 | 114.701 | -0.05774 | | 1 | 400 | 6460 | 6.46 | -0.14434 |

| | | | | | | | | | | |
|---|---|---|---|---|---|---|---|---|---|---|
| 2 | 500 | 94291 | 94.291 | -0.05340 | | 2 | 500 | 5264 | 5.264 | -0.14520 |
| 3 | 600 | 76516 | 76.516 | -0.04678 | | 3 | 600 | 4249 | 4.249 | -0.14544 |
| 4 | 700 | 60606 | 60.606 | -0.03795 | | 4 | 700 | 3301 | 3.301 | -0.14855 |
| 5 | 800 | 45533 | 45.533 | -0.02689 | | 5 | 800 | 2435 | 2.435 | -0.14170 |
| 6 | 900 | 30369 | 30.369 | -0.01515 | -0.03645 | 6 | 900 | 1588 | 1.588 | -0.12991 |

| | Table S7: Previously published divergence time estimates between *Pavo cristatus* and *Pavo muticus*. | | | | |
|---|---|---|---|---|---|
| **Sr. No.** | **Title** | **Author** | **Journal** | **Year** | **Split time (MYA)** |
| 1 | Resolution of the Phylogenetic Position of the Congo Peafowl, | Kimball et al. | Proc R Soc B | 1997 | |
| | Afropavo congensis: A Biogeographic and Evolutionary Enigma | | | | |
| | cytochrome b - 4.2 mya | | | | 4.2 |
| | D-loop - 6.2 mya | | | | 6.2 |
| 2 | The global diversity of birds in space and time | Jetz et al. | Nature | 2012 | |
| | nuclear + mitochondrial markers tree - 1.5 Mya | | | | 1.5 |
| 3 | A molecular genetic time scale demonstrates Cretaceous origins and | Stein et al. | MPE | 2015 | 2.27 |
| | multiple diversification rate shifts within the order Galliformes (Aves) | | | | |
| | nuclear + mitochondrial markers tree | | | | |
| 4 | Genetic Divergence between Pavo muticus and Pavo cristatus | Ouyang, Y. N., et al. | Journal of Yunnan | 2009 | 2.96 |
| | by Cyt b gene. | | Agricultural University | | |
| | Cyt b | | | | |
| 5 | A Phylogenomic Supertree of Birds | Kimball et al. | Diversity | 2019 | 3.17 |
| | WGS+UCE supertree | | | | |
| 6 | Ancestral range reconstruction of Galliformes: the effects of topology | Wang et al. | Journal of Biogeography | 2016 | 2.58 |
| | and taxon sampling | | | | |
| | nuclear + mitochondrial markers tree | | | | |
| 7 | Divergence time estimation of Galliformes based on the best gene | Chen et al. | BMC Ecology and | 2021 | 3.12 |
| | shopping scheme of ultraconserved elements | | Evolution | | |
| | UCE | | | | |
| 8 | this study (UCE) | | | | 2.02 |
| 9 | this study (PSMC) | | | | 1.1 |

**Table S8: Amino acid changes across KEGG signalling pathways.**

| | | | | |
|---|---|---|---|---|
| **1. Hedgehog** | | | | |
| **Sr No** | **Gene** | **position** | ***P. muticus*** | ***P. cristatus*** |
| **1** | EFCAB7 | 407 | L | S |
| **2** | HHAT | 205 | R | Q |
| **3** | KIF7 | 695 | L | S |
| | KIF7 | 1292 | T | A |
| **4** | MGRN1 | 504 | T | S |
| **2. Melanogenesis** | | | | |
| **1** | CREB3 | 339 | A | T |
| **2** | PLCB1 | 908 | L | S |
| **3. TGF-beta** | | | | |
| **1** | RBL1 | 211 | F | L |
| | RBL1 | 507 | L | S |
| **2** | THBS1 | 797 | N | S |
| **3** | ZFYVE16 | 31 | A | T |
| | ZFYVE16 | 282 | C | Y |
| | ZFYVE16 | 335 | S | F |
| | ZFYVE16 | 599 | Q | E |
| **4. Wnt** | | | | |
| **1** | APC | 277 | G | A |
| | APC | 877 | P | A |
| | APC | 909 | M | T |
| | APC | 1216 | P | Q |
| | APC | 1309 | D | H |
| **2** | GPC4 | 556 | V | M |
| **3** | INVS | 828 | I | R |
| **4** | PLCB1 | 908 | L | S |
| **5** | ROR2 | 149 | H | Y |
| **5. Notch** | | | | |
| **1** | DLL1 | 573 | I | V |
| **2** | DLL4 | 385 | I | V |
| **6. Extracellular matrix interaction** | | | | |
| **1** | AGRN | 786 | G | S |
| **2** | COL4A2 | 362 | V | L |
| **3** | GP1BB | 142 | A | T |
| | GP1BB | 179 | A | T |
| **4** | HSPG2 | 966 | R | H |

| | | | | |
|---|---|---|---|---|
| **5** | ITGB1 | 12 | V | I |
| **6** | LAMA4 | 435 | H | R |
| **7** | LAMA5 | 686 | M | T |
| | LAMA5 | 1728 | V | I |
| | LAMA5 | 1814 | A | G |
| | LAMA5 | 3225 | W | R |
| | LAMA5 | 3450 | I | V |
| | LAMA5 | 3560 | T | M |
| **8** | LAMC1 | 272 | A | S |
| | LAMC1 | 1075 | V | I |
| | LAMC1 | 1208 | G | D |
| **9** | SDC1 | 180 | P | L |
| **10** | THBS1 | 797 | N | S |
| **11** | VWF | 652 | G | S |
| **7. Actin cytoskeleton** | | | | |
| **1** | ACTN1 | 851 | E | D |
| **2** | ACTN4 | 850 | E | D |
| **3** | APC | 277 | G | A |
| | APC | 877 | P | A |
| | APC | 909 | M | T |
| | APC | 1216 | P | Q |
| | APC | 1309 | D | H |
| **5** | ITGB1 | 12 | V | I |
| **6** | MYLK | 920 | G | V |
| **7** | NCKAP1L | 579 | I | V |
| **8** | SSH2 | 1218 | C | G |
| **9** | SSH3 | 409 | Q | R |
| **10** | VCL | 534 | M | L |
| **8. Focal adhesion** | | | | |
| **1** | KDR | 1336 | V | A |
| **2** | MYLK | 920 | G | V |
| **3** | VEGFC | 265 | T | A |
| **4** | ZYX | 289 | V | A |
| **5** | ITGB1 | 12 | V | I |
| **6** | VCL | 534 | M | L |
| **7** | ACTN1 | 851 | E | D |
| **8** | ACTN4 | 850 | E | D |
| **9** | VWF | 652 | G | S |
| **10** | THBS1 | 797 | N | S |

| | | | | |
|---|---|---|---|---|
| **11** | LAMA4 | 435 | H | R |
| **12** | LAMA5 | 686 | M | T |
| | LAMA5 | 1728 | V | I |
| | LAMA5 | 1814 | A | G |
| | LAMA5 | 3225 | W | R |
| | LAMA5 | 3450 | I | V |
| | LAMA5 | 3560 | T | M |
| **13** | LAMC1 | 272 | A | S |
| | LAMC1 | 1075 | V | I |
| | LAMC1 | 1208 | G | D |
| **14** | COL4A2 | 362 | V | L |
| **9. MAPK** | | | | |
| 1 | ATF4 | 257 | Q | R |
| 2 | CHUK | 743 | K | R |
| 3 | CSF1R | 405 | H | R |
| 4 | IGF2 | 122 | S | N |
| 5 | IKBKB | 701 | V | M |
| 6 | INSR | 995 | T | A |
| | INSR | 998 | H | R |
| 7 | MAP3K13 | 929 | A | V |
| 8 | MAP3K4 | 76 | V | I |
| 9 | MAP4K3 | 299 | T | S |
| 10 | NF1 | 544 | E | D |
| 11 | NFKB2 | 513 | Q | R |
| 12 | PLA2G4A | 153 | G | E |
| 13 | PLA2G4EL6 | 478 | V | I |
| 14 | PLA2G4F | 596 | K | R |
| 15 | TRAF6 | 45 | L | P |

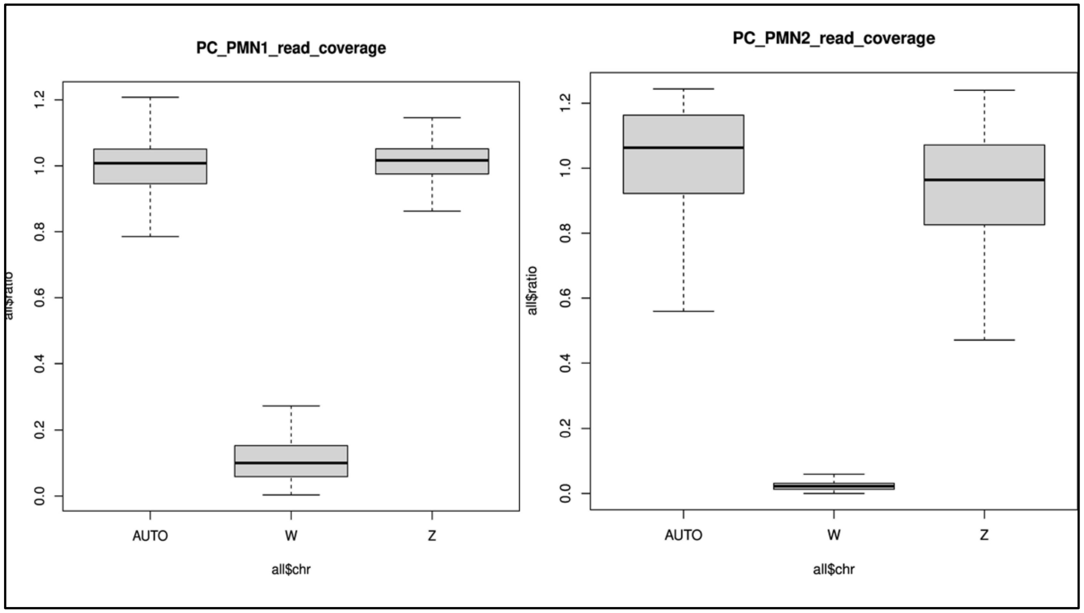

**Figure S1: Mean read coverage of autosomes, W chromosome and Z chromosome of *P. cristatus* individual 1 (left) and individual 2 (right). Equivalent read coverages of autosomes and Z chromosome (~1), whereas negligible for W chromosome (<0.1) shows characteristics of male karyotype.**

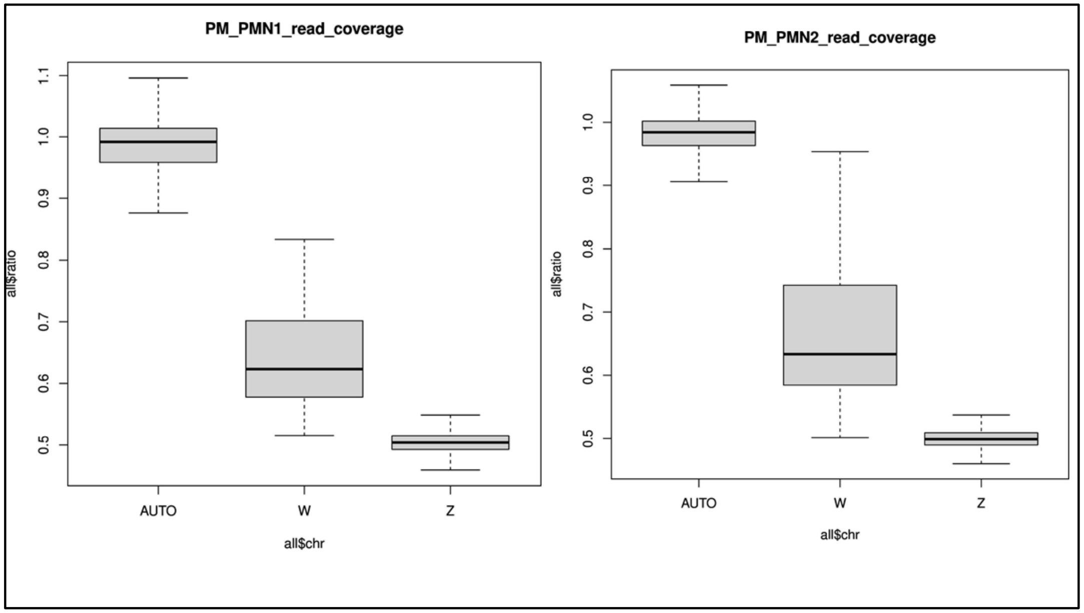

**Figure S2: Mean read coverage of autosomes, W chromosome and Z chromosome of *P. muticus* individual 1 (left) and individual 2 (right). Read coverage for autosomes is ~1 and for Z and W chromosome ~0.5, are characteristics of a female karyotype.**

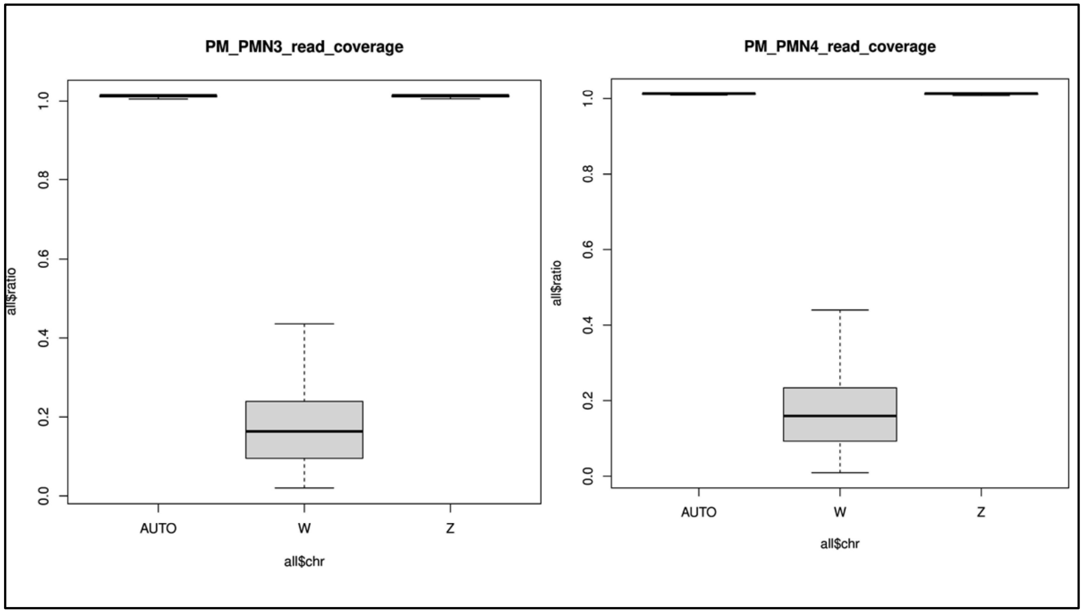

**Figure S3: Mean read coverage of autosomes, W chromosome and Z chromosome of *P. muticus* individual 3 (left) and individual 4 (right). Equivalent read coverages of autosomes and Z chromosome (~1), whereas negligible for W chromosome (~0.1) shows characteristics of male karyotype.**

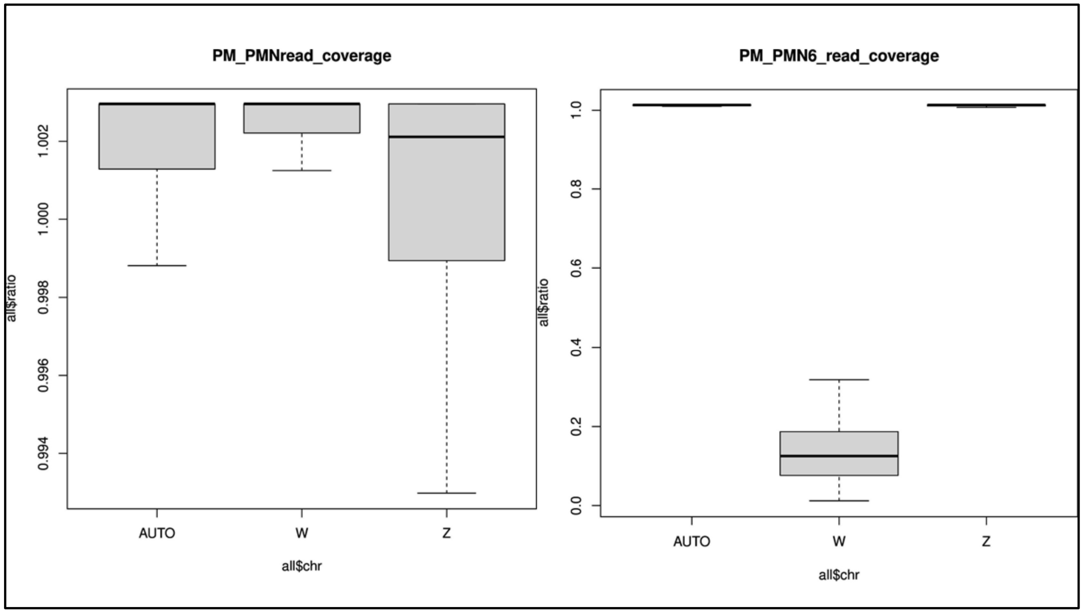

**Figure S4: Mean read coverage of autosomes, W chromosome and Z chromosome of *P. muticus* individual 5 (left) and individual 6 (right). Equivalent read coverages of autosomes and Z chromosome (~1), whereas negligible for W chromosome (<0.1) shows characteristics of male karyotype.**

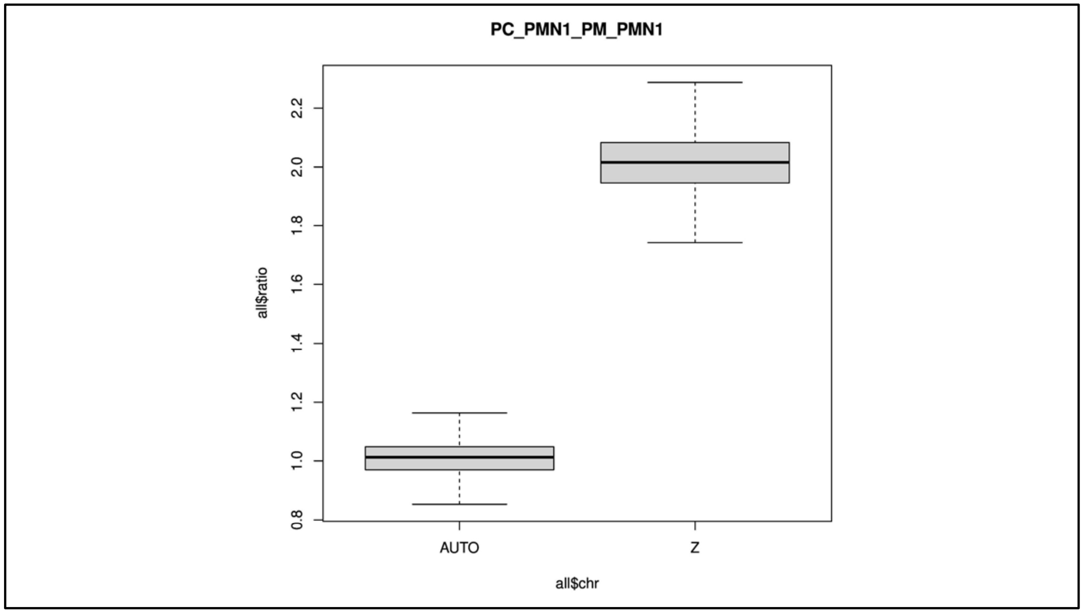

**Figure S5: Mean read coverage ratio of autosomes and Z chromosome of *P. cristatus* individual 1 (male) and *P. muticus* individual 1. The ratio of read coverage of autosomes is 1 whereas of Z chromosome is 2, which shows that *P. muticus* individual 1 is female.**

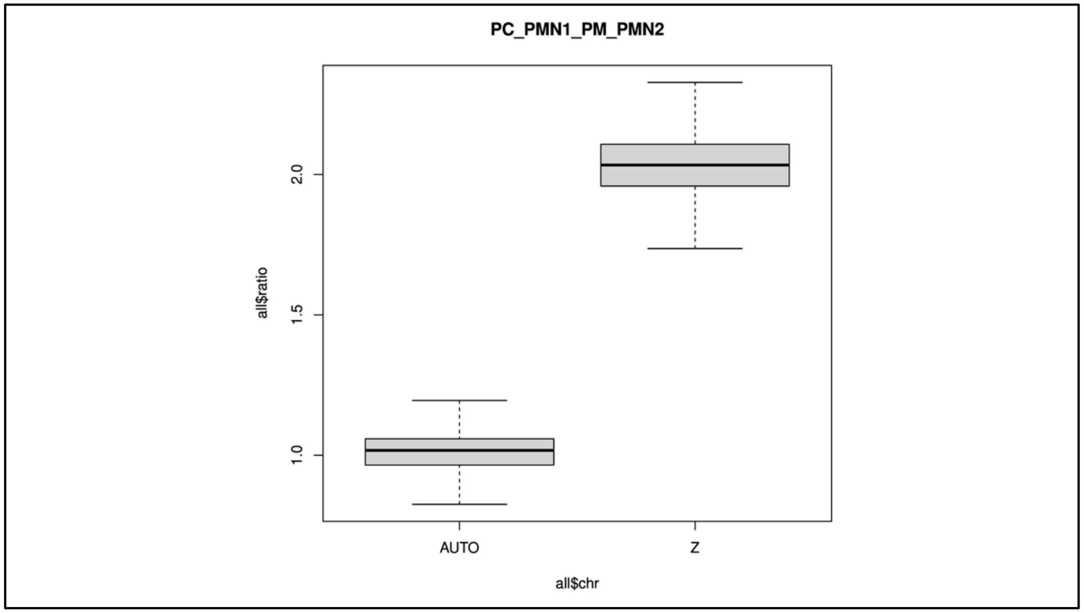

**Figure S6: Mean read coverage ratio of autosomes and Z chromosome of *P. cristatus* individual 1 (male) and *P. muticus* individual 2. The ratio of read coverage of autosomes is 1 whereas of Z chromosome is 2, which shows that *P. muticus* individual 2 is female.**

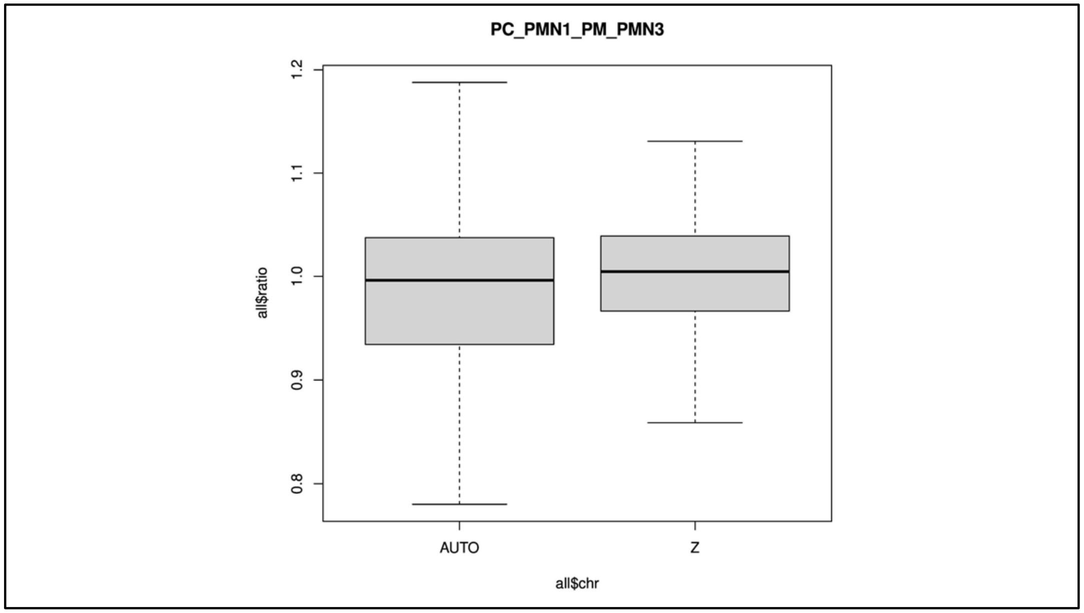

**Figure S7: Mean read coverage ratio of autosomes and Z chromosome of *P. cristatus* individual 1 (male) and *P. muticus* individual 3. The ratio of read coverage of autosomes and Z chromosome is 1, which shows that *P. muticus* individual 3 is male.**

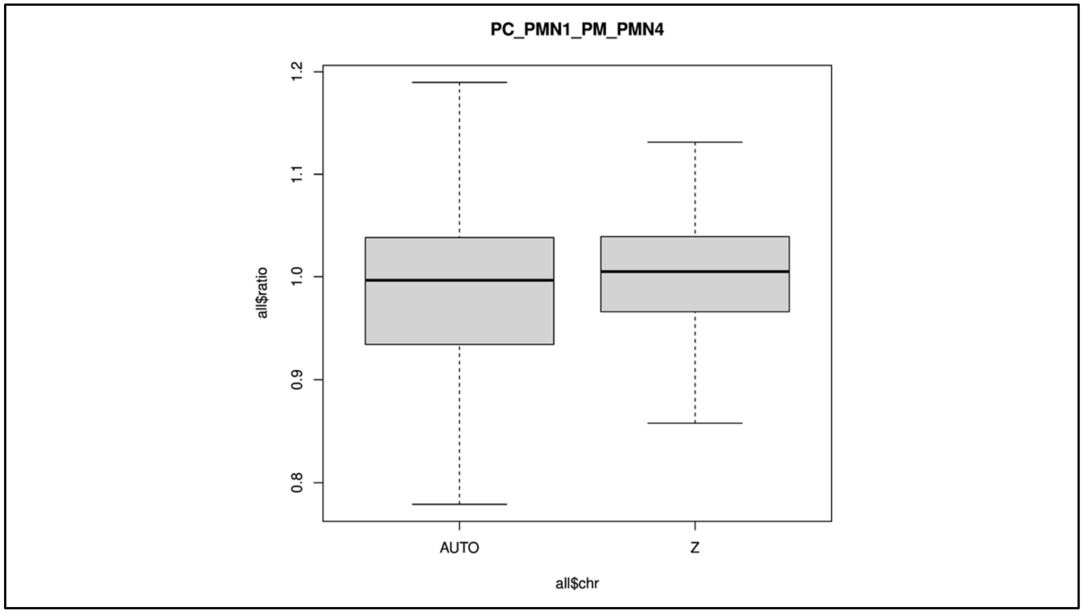

**Figure S8: Mean read coverage ratio of autosomes and Z chromosome of *P. cristatus* individual 1 (male) and *P. muticus* individual 4. The ratio of read coverage of autosomes and Z chromosome is 1, which shows that *P. muticus* individual 4 is male.**

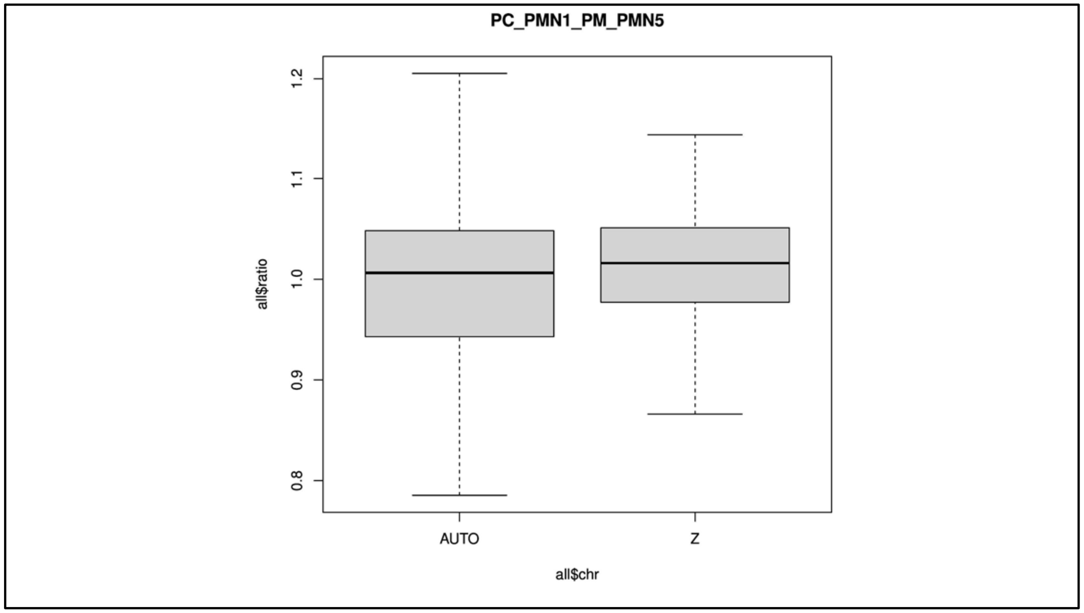

**Figure S9: Mean read coverage ratio of autosomes and Z chromosome of *P. cristatus* individual 1 (male) and *P. muticus* individual 5. The ratio of read coverage of autosomes and Z chromosome is 1, which shows that *P. muticus* individual 5 is male.**

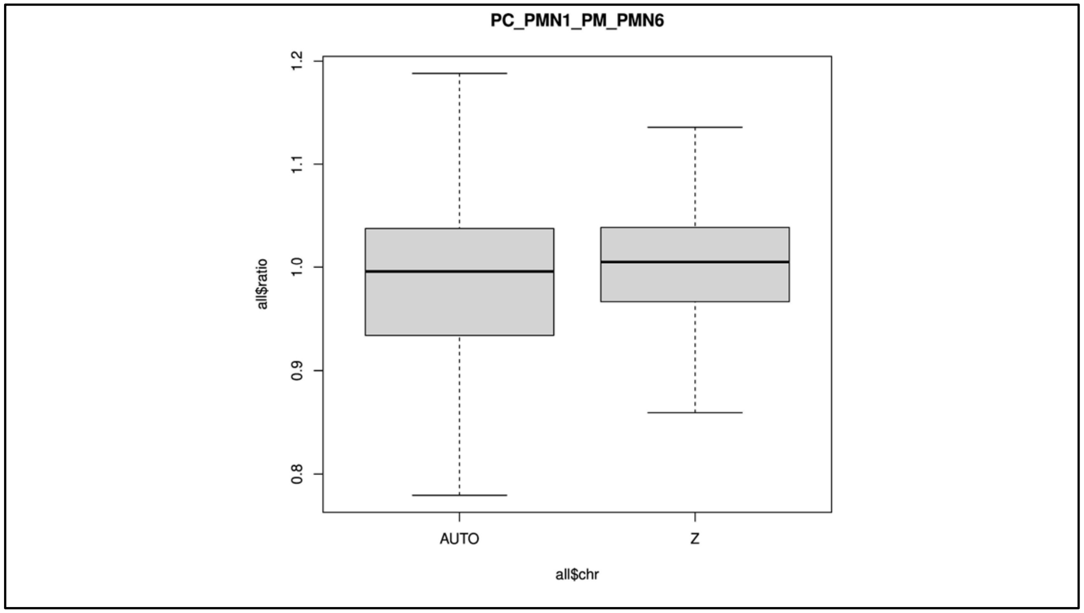

**Figure S10: Mean read coverage ratio of autosomes and Z chromosome of *P. cristatus* individual 1 (male) and *P. muticus* individual 6. The ratio of read coverage of autosomes and Z chromosome is 1, which shows that *P. muticus* individual 6 is male.**

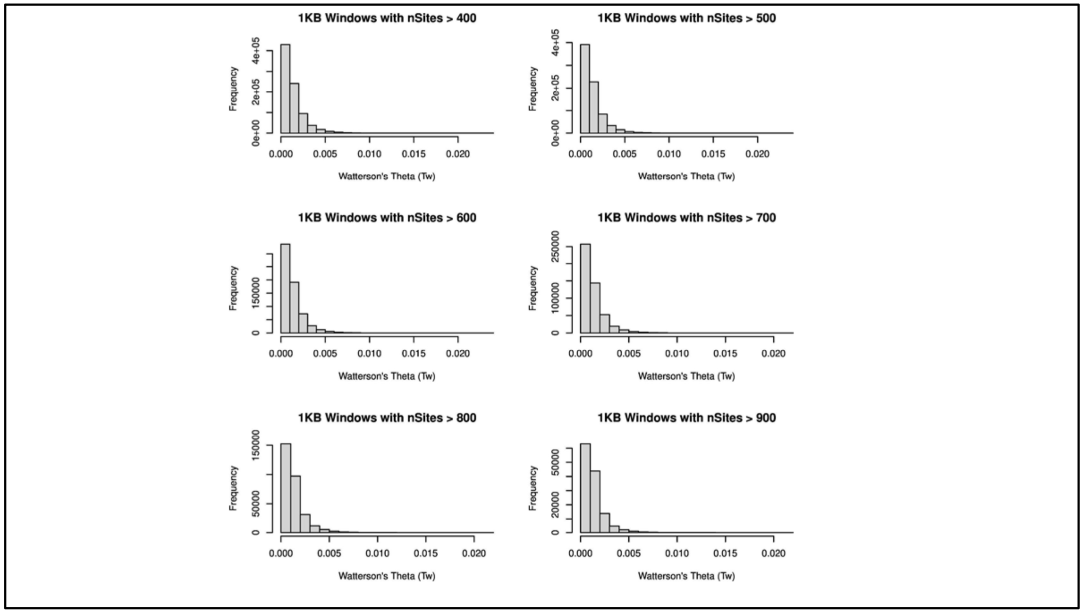

**Figure S11: Histograms of Watterson's theta distributions of autosomes of *P. muticus* with cutoffs of nsites from greater than 400 to 900.**

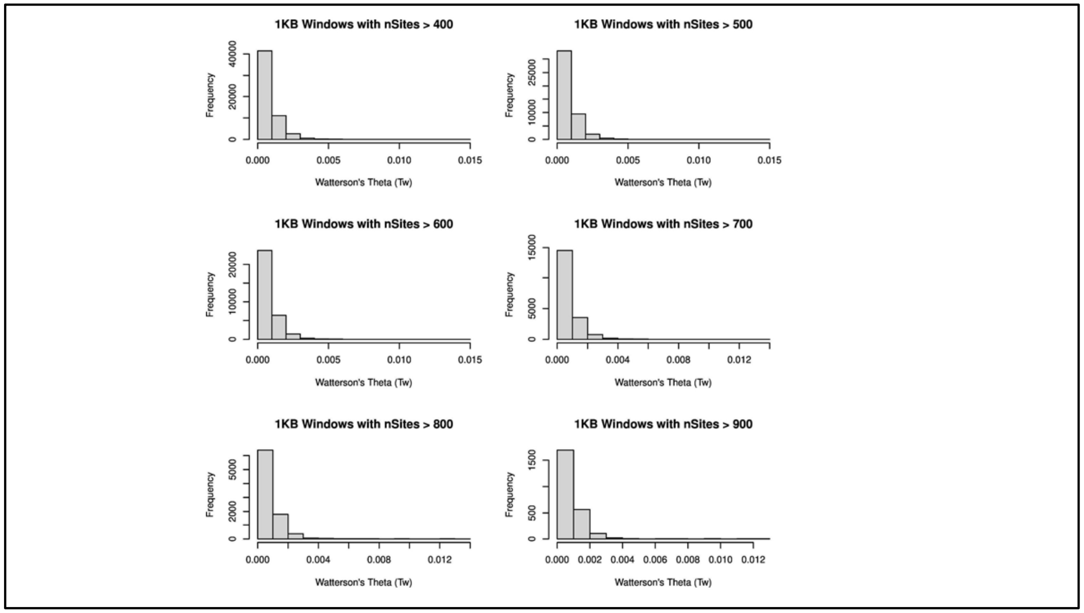

**Figure S12: Histograms of Watterson's theta distributions of Z chromosome of *P. muticus* with cutoffs of nsites from greater than 400 to 900.**

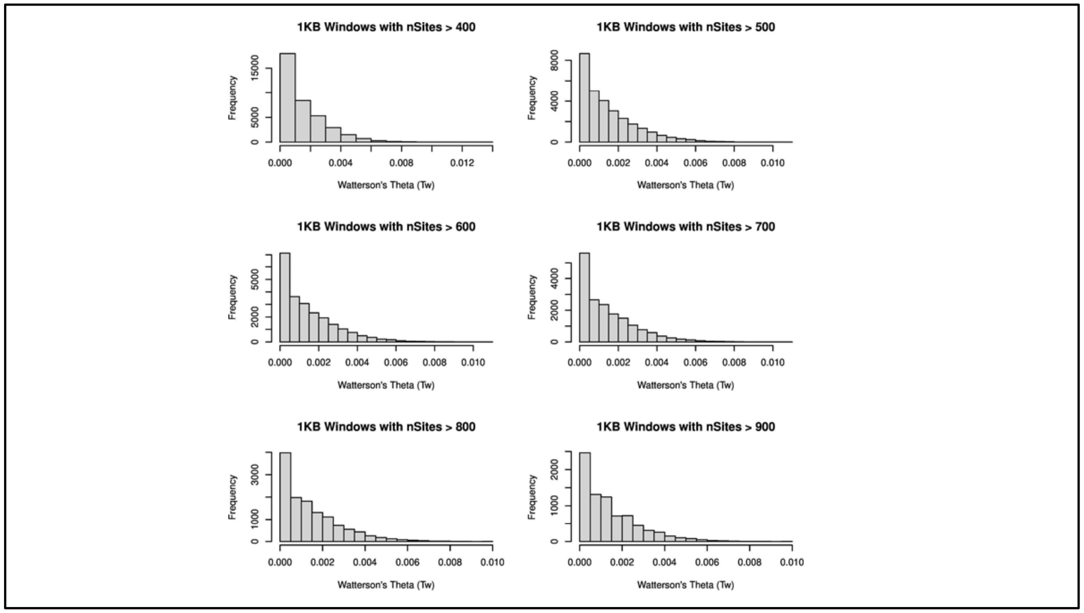

**Figure S13: Histograms of Watterson's theta distributions of autosomes of *P. cristatus* with cutoffs of nsites from greater than 400 to 900.**

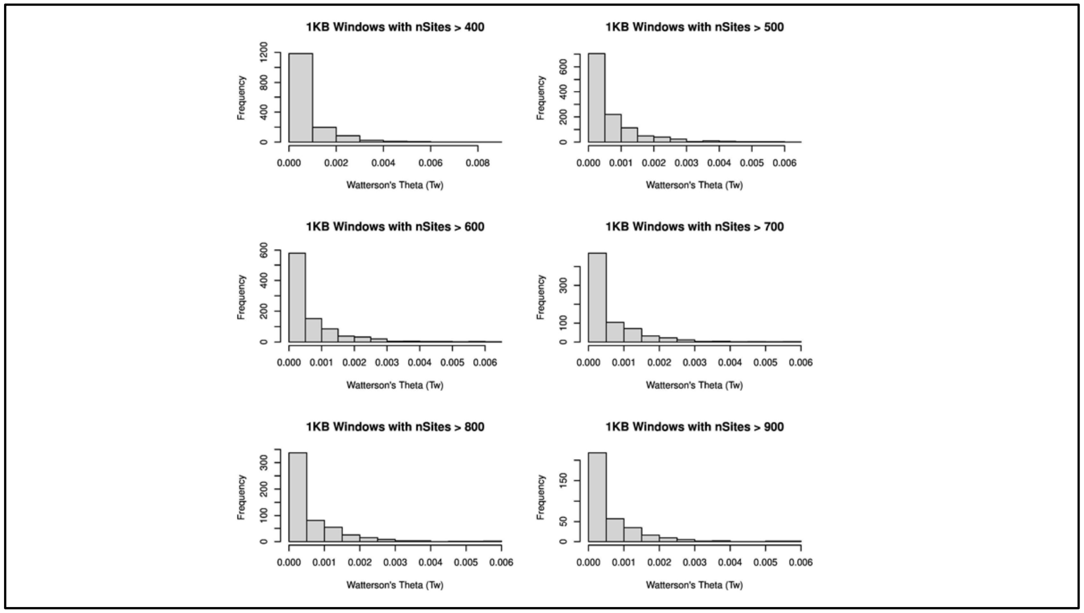

**Figure S14: Histograms of Watterson's theta distributions of Z chromosome of *P. cristatus* with cutoffs of nsites from greater than 400 to 900.**

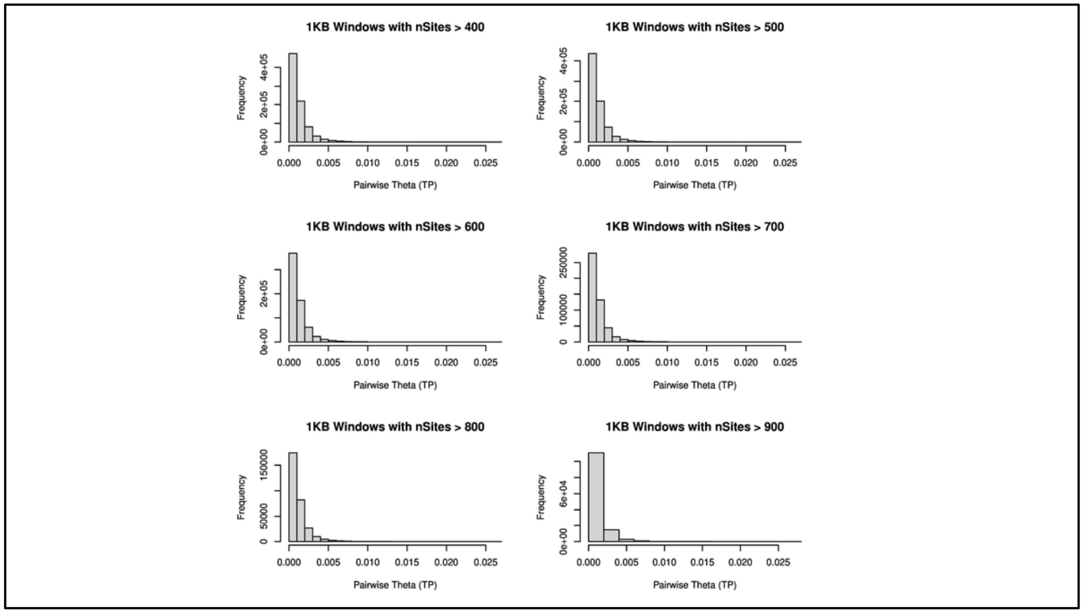

**Figure S15: Histograms of  pairwise theta distributions of autosomes of *P. muticus* with cutoffs of nsites from greater than 400 to 900.**

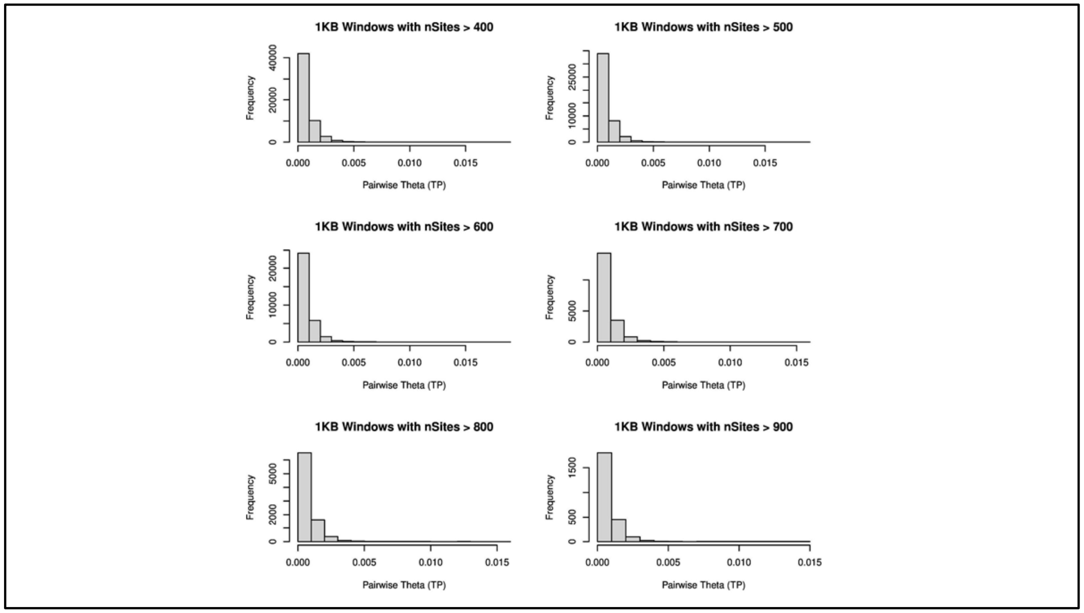

**Figure S16: Histograms of pairwise theta distributions of Z chromosome of *P. muticus* with cutoffs of nsites from greater than 400 to 900.**

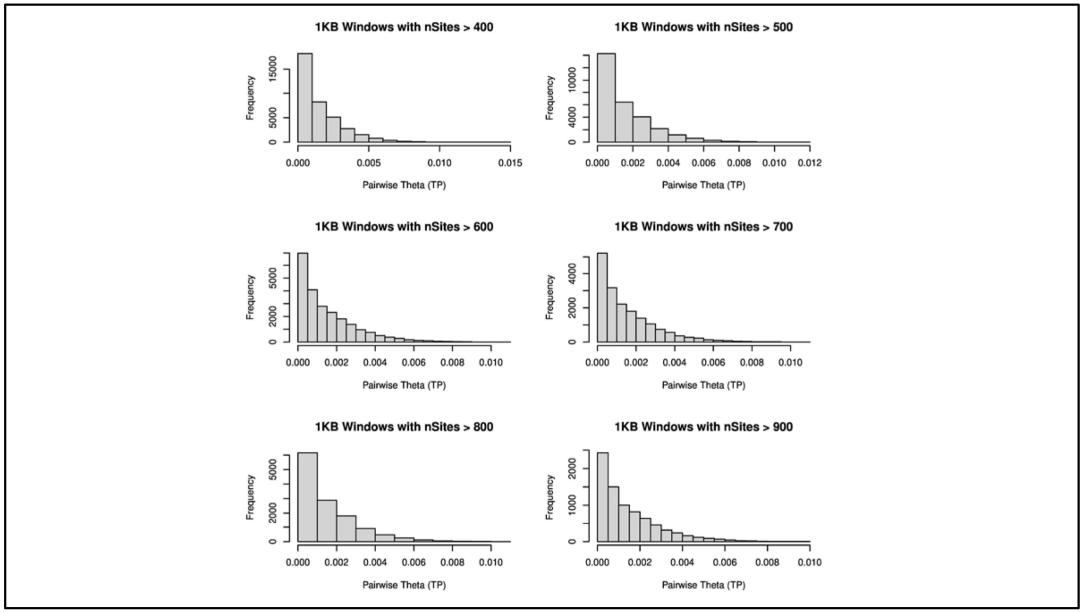

**Figure S17: Histograms of pairwise theta distributions of autosomes of *P. cristatus* with cutoffs of nsites from greater than 400 to 900.**

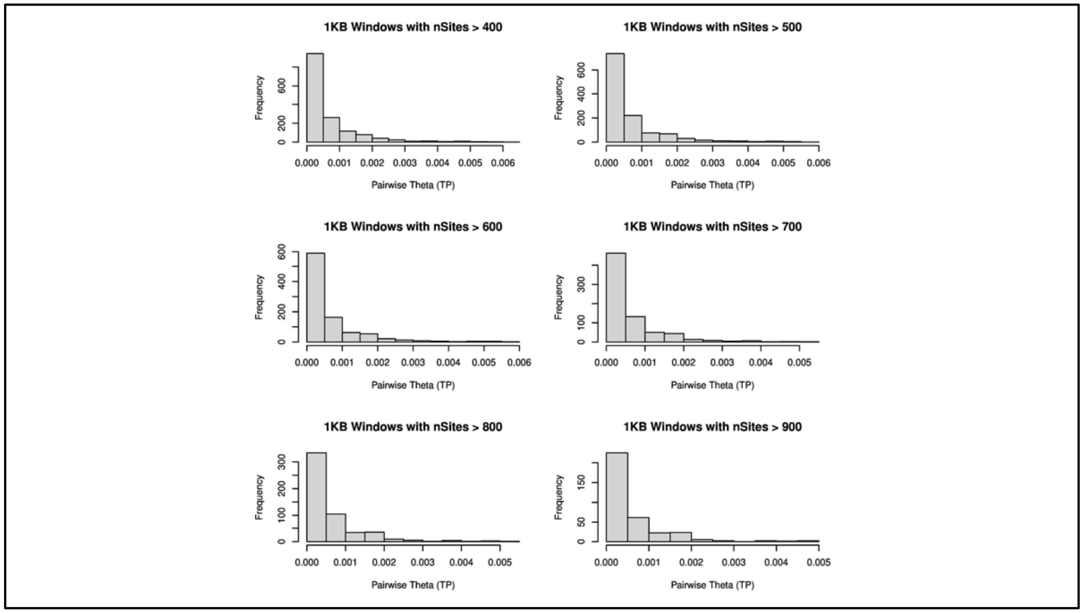

**Figure S18: Histograms of pairwise theta distributions of Z chromosome of *P. cristatus* with cutoffs of nsites from greater than 400 to 900.**

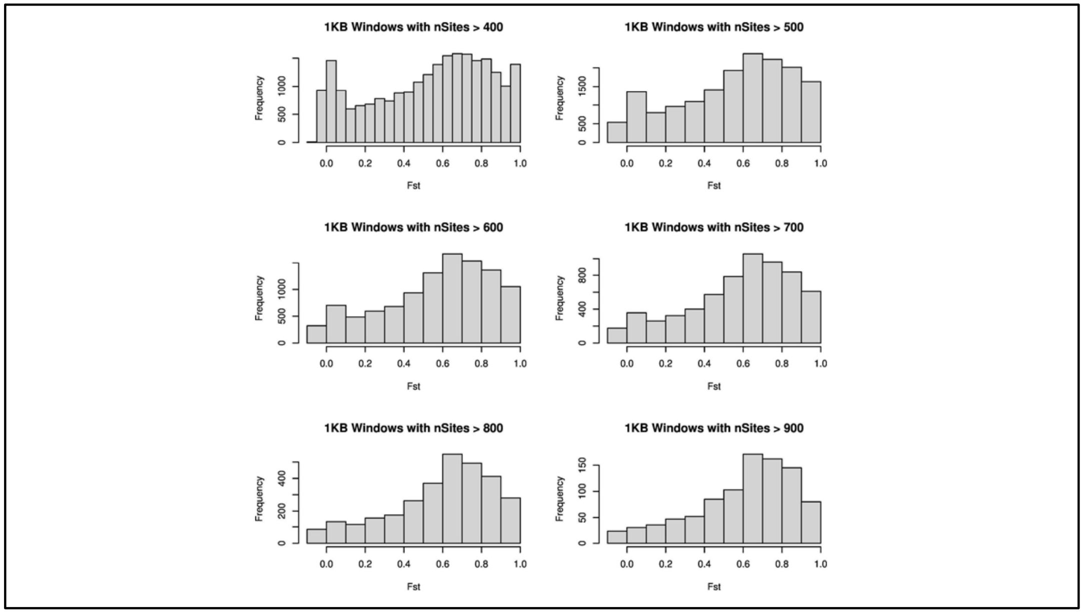

**Figure S19: Histograms of FST distributions of autosomes with cutoffs of nsites from greater than 400 to 900.**

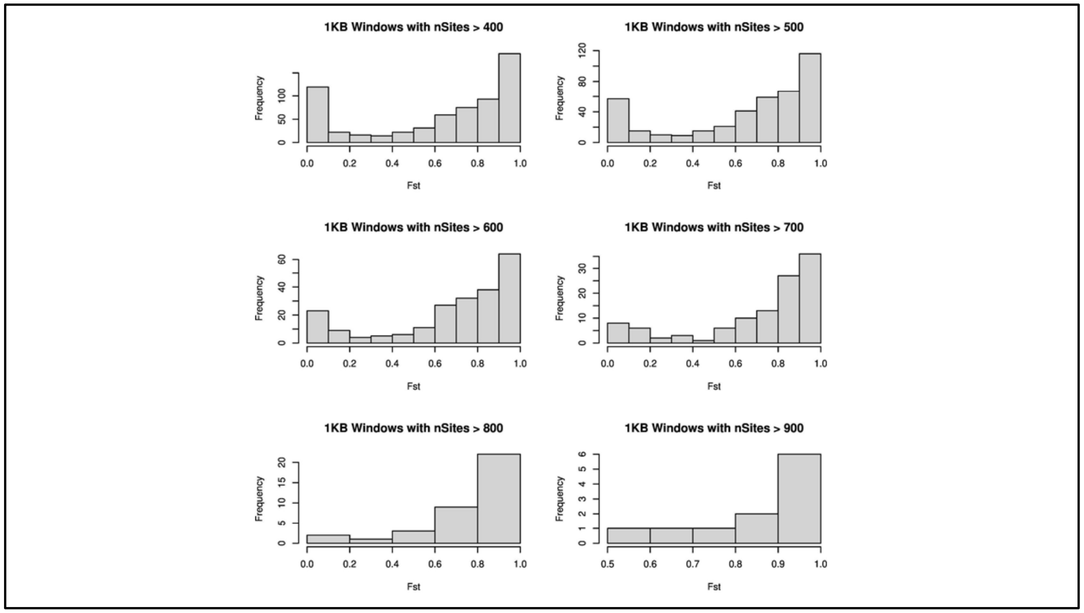

**Figure S20: Histograms of FST distributions of Z chromosome with cutoffs of nsites from greater than 400 to 900.**

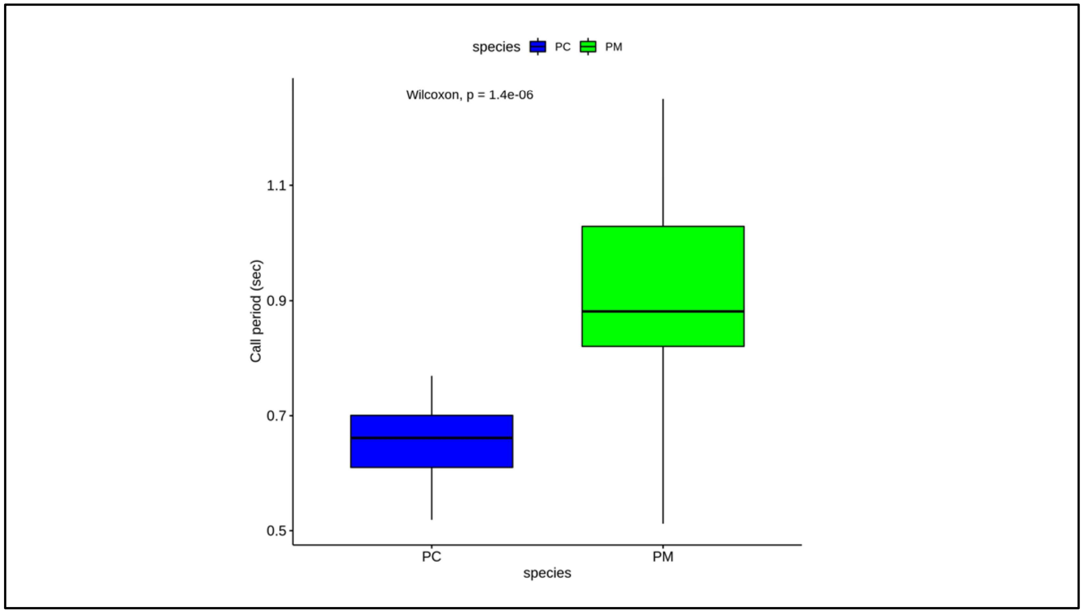

**Figure S21: Temporal acoustic property (Delta time) comparison of *Pavo cristatus* (blue) and *Pavo muticus* (green) calls.**
Boxplots comparing call periods of two species. *Pavo cristatus* (blue) has a significantly shorter call than *Pavo muticus* (green) (Pairwise Wilcoxon test, $p < 0.0001$).

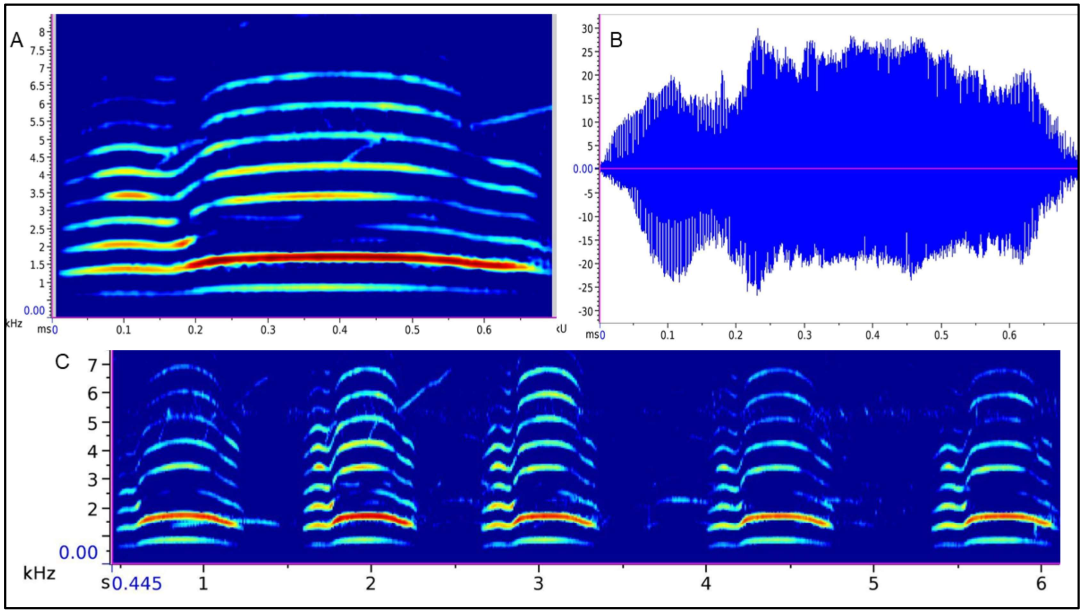

**Figure S22: Visualisations of acoustic properties recorded of *Pavo cristatus*.**

A. The spectrogram view of a single call of *Pavo cristatus*.
B. The waveform view of a single call of *Pavo cristatus*.
C. The spectrogram view of a call group of *Pavo cristatus*.

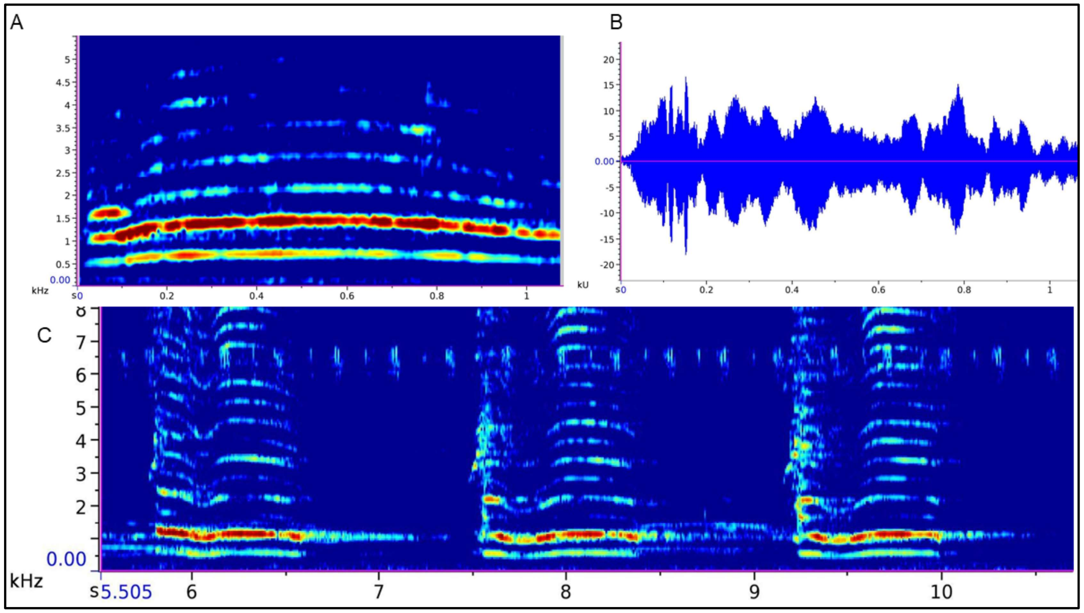

**Figure S23: Visualisations of acoustic properties recorded of *Pavo muticus*.**

A. The spectrogram view of a single call of *Pavo muticus*.
B. The waveform view of a single call of *Pavo muticus*.
C. The spectrogram view of a call group of *Pavo muticus*.

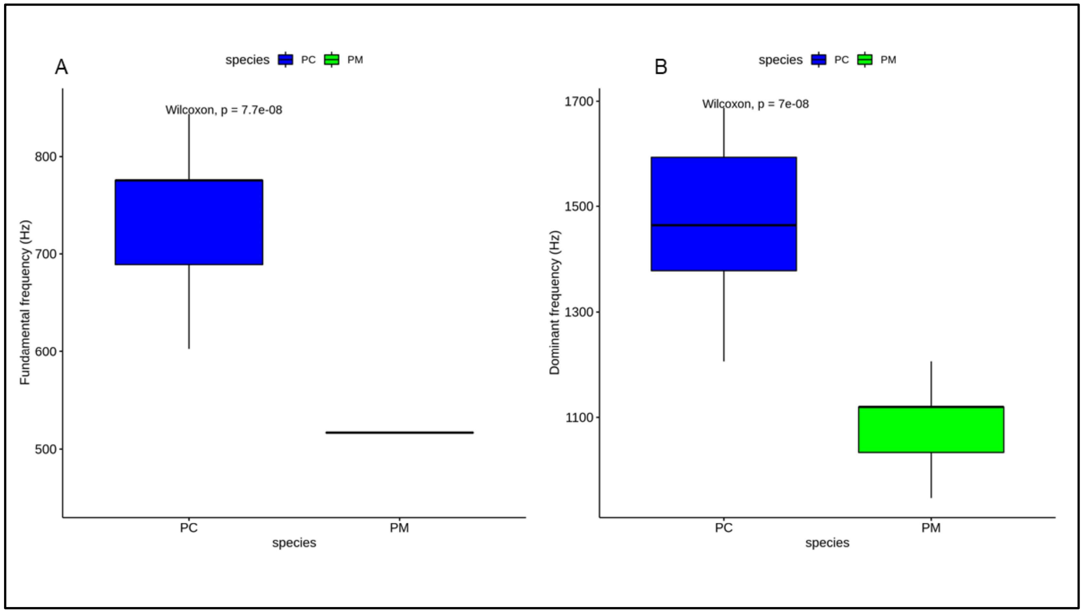

**Figure S24: Spectral properties of calls recorded in this study.**

*A.* *Pavo cristatus* (blue) has a significantly higher fundamental frequency than *Pavo muticus* (green) (Pairwise Wilcoxon test, $p < 0.0001$).

*B.* *Pavo cristatus* (blue) has a significantly higher dominant frequency than *Pavo muticus* (green) (Pairwise Wilcoxon test, $p < 0.0001$).

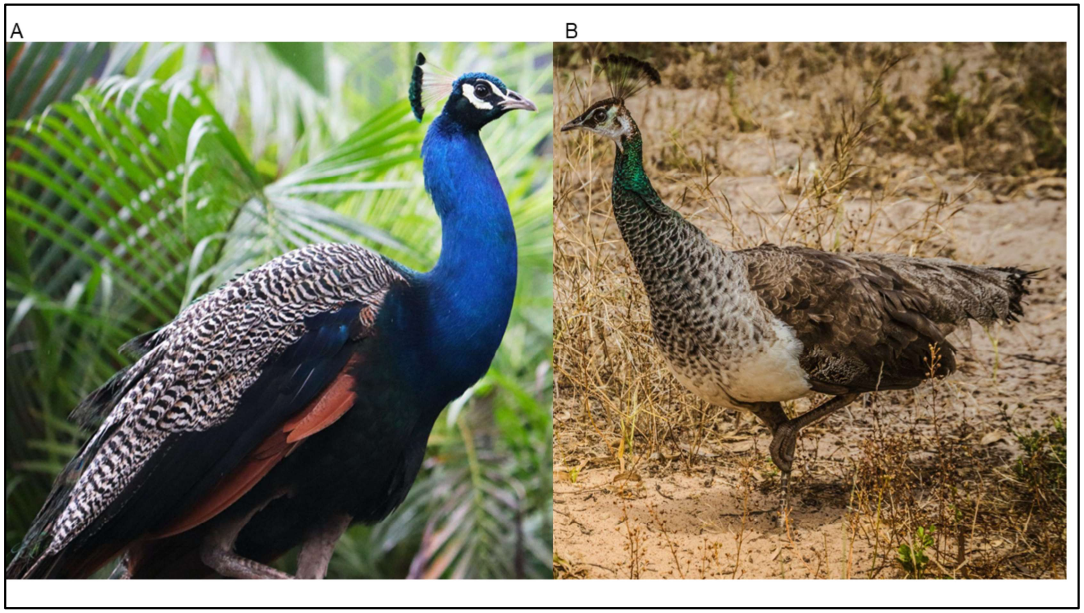

**Figure S25: The sexual diamorphism in *Pavo cristatus*.**

A. The male *Pavo cristatus*.
B. The female *Pavo cristatus*.

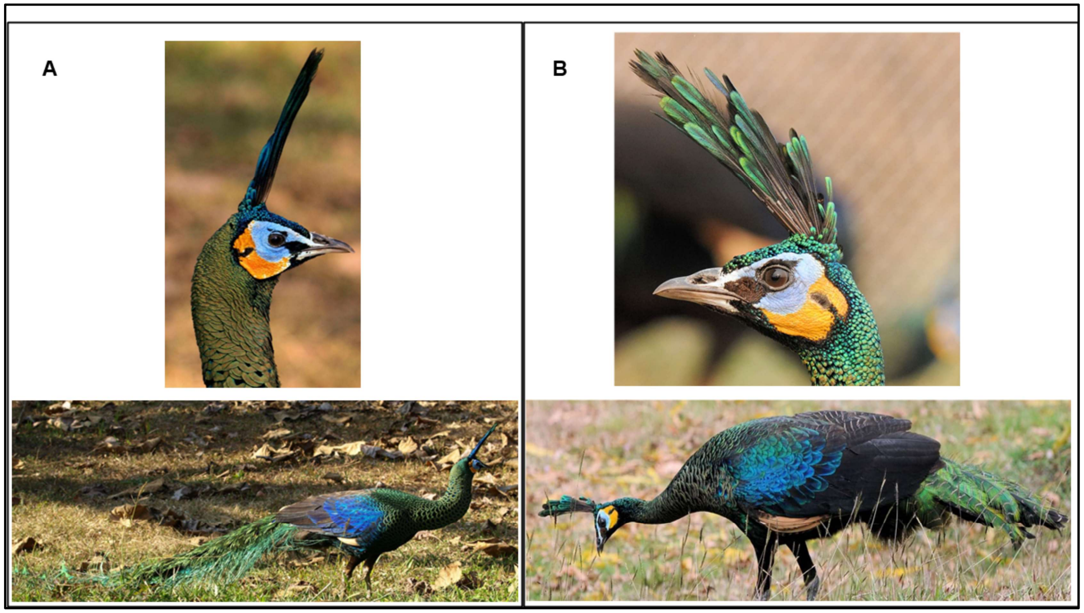

**Figure S26: The sexual diamorphism in *Pavo muticus.***

A. The male *Pavo muticus.*
B. The female *Pavo muticus*.

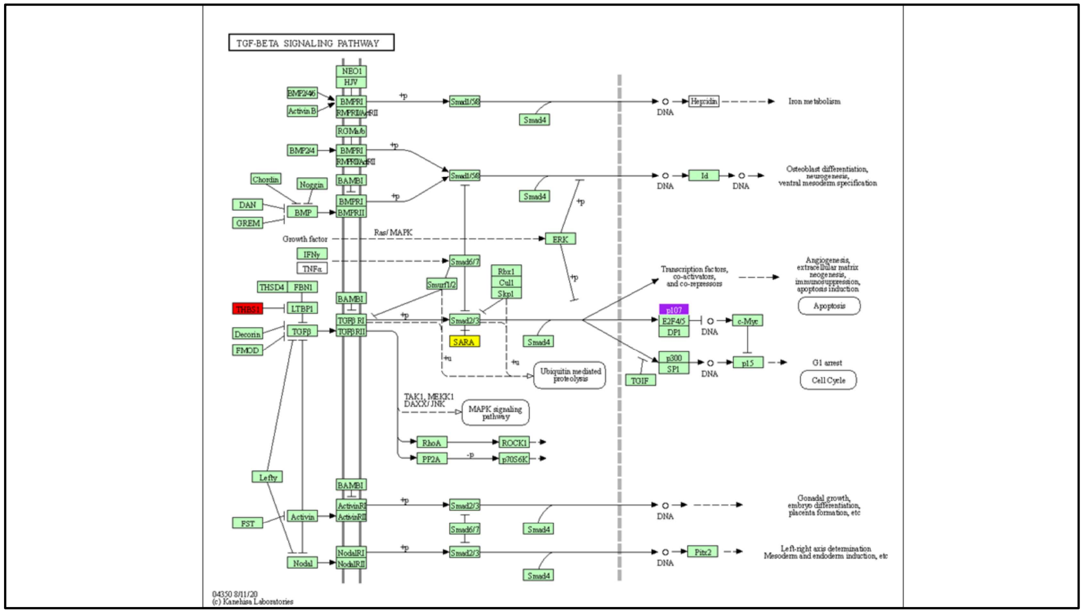

**Figure S27: The KEGG annotation of the TGF-Beta signalling pathway.**

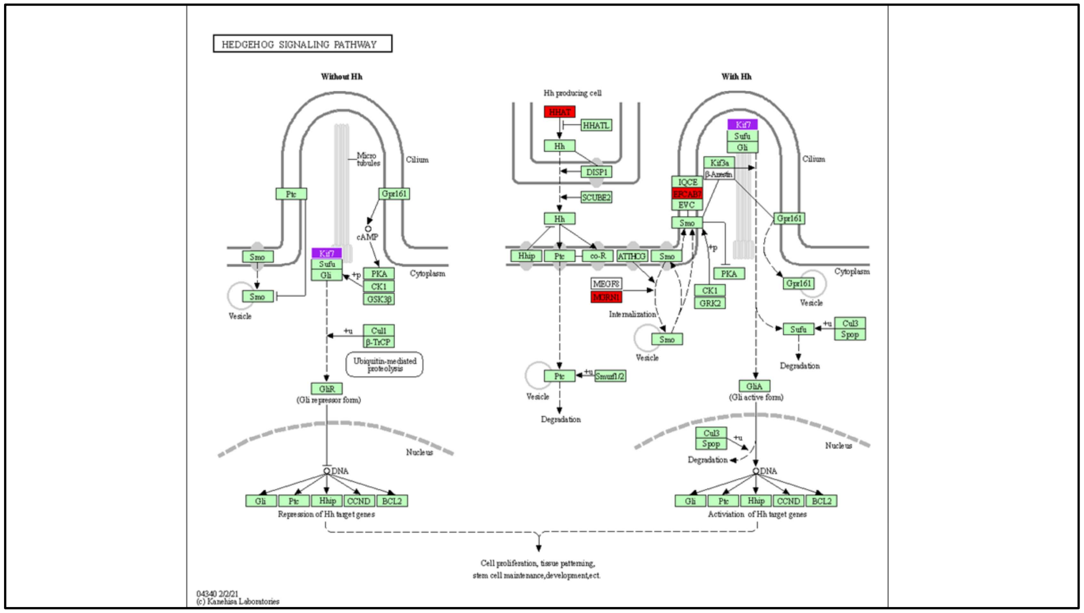

**Figure S28: The KEGG annotation of the Hedgehog signalling pathway.**

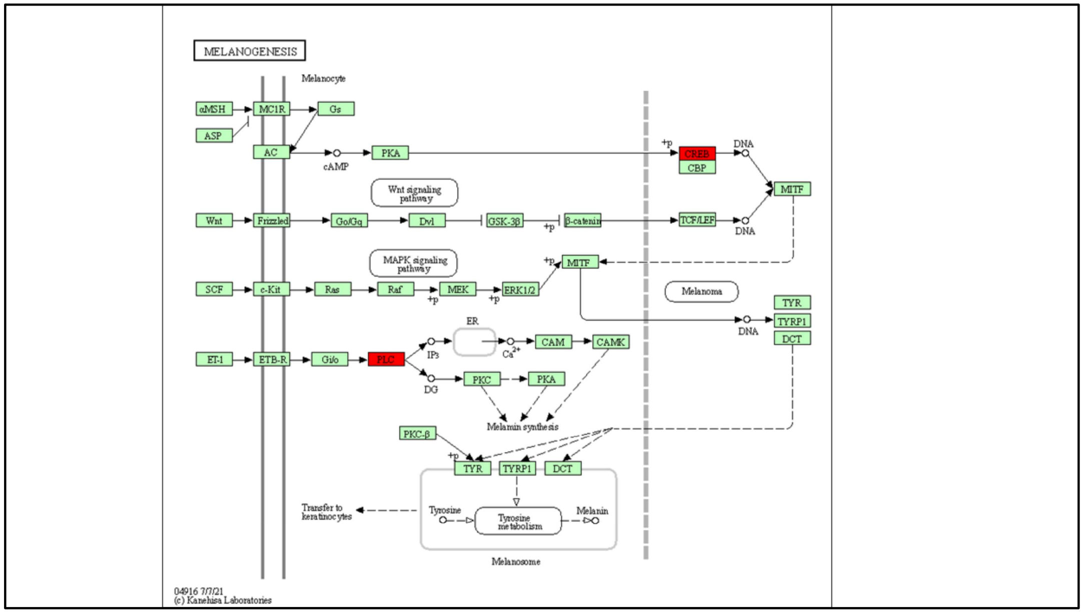

**Figure S29: The KEGG annotation of the Melanogenesis pathway.**

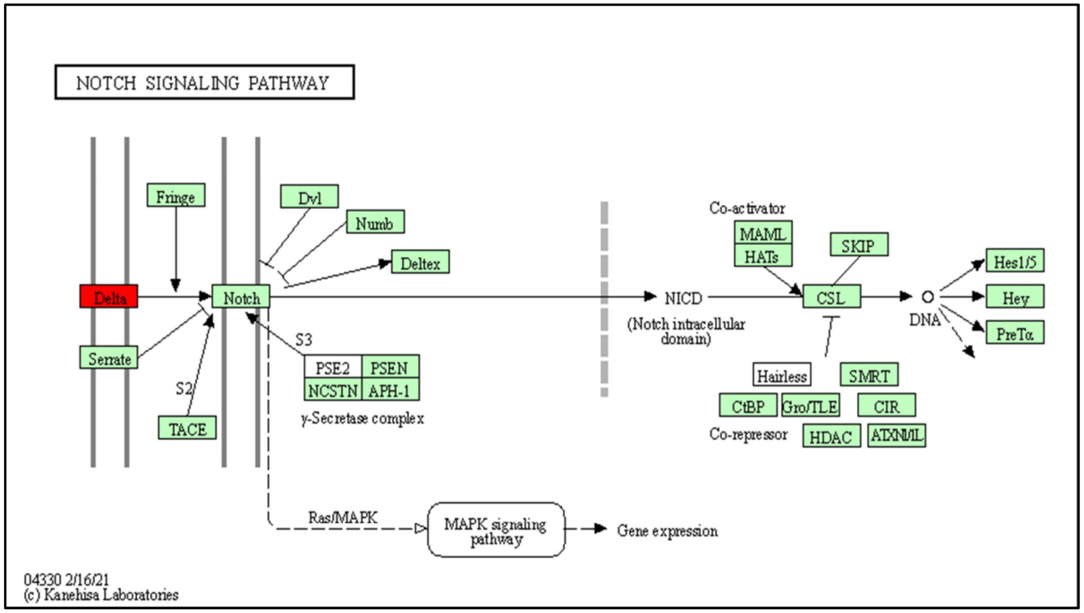

**Figure S30: The KEGG annotation of the Notch signalling pathway.**

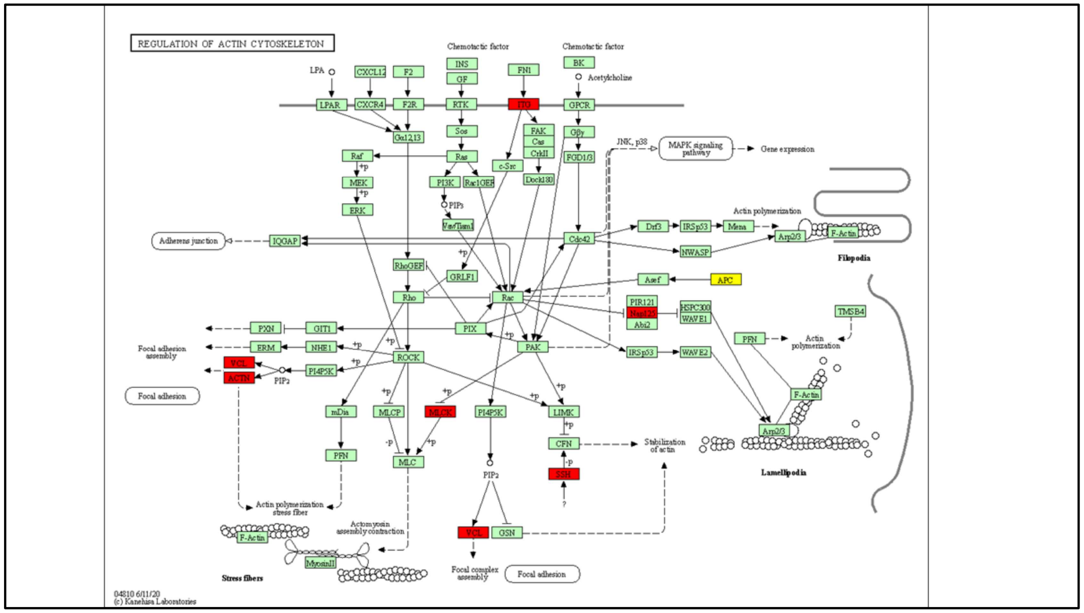

**Figure S31: The KEGG annotation of the Regulation of actin cytoskeleton pathway.**

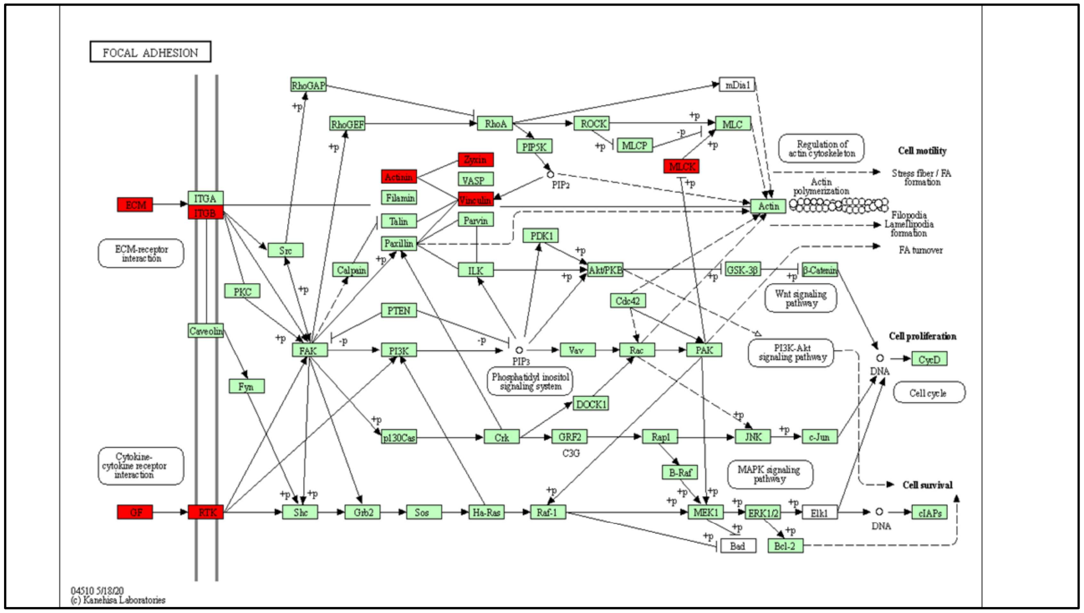

**Figure S32: The KEGG annotation of the focal adhesion pathway.**

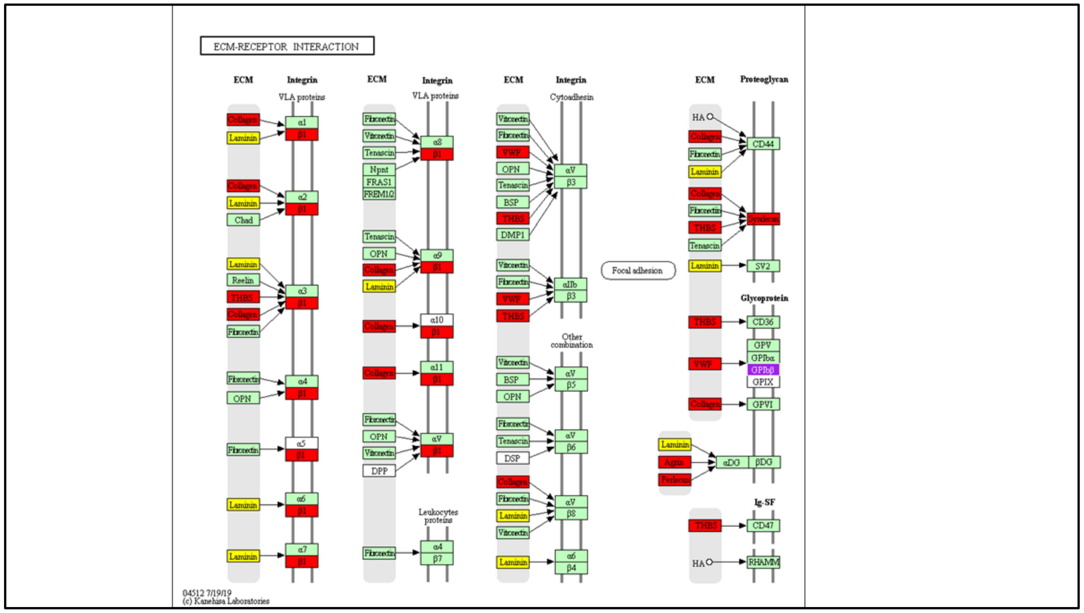

**Figure S33: The KEGG annotation of the Extracellular matrix (ECM) receptor interaction signalling pathway.**

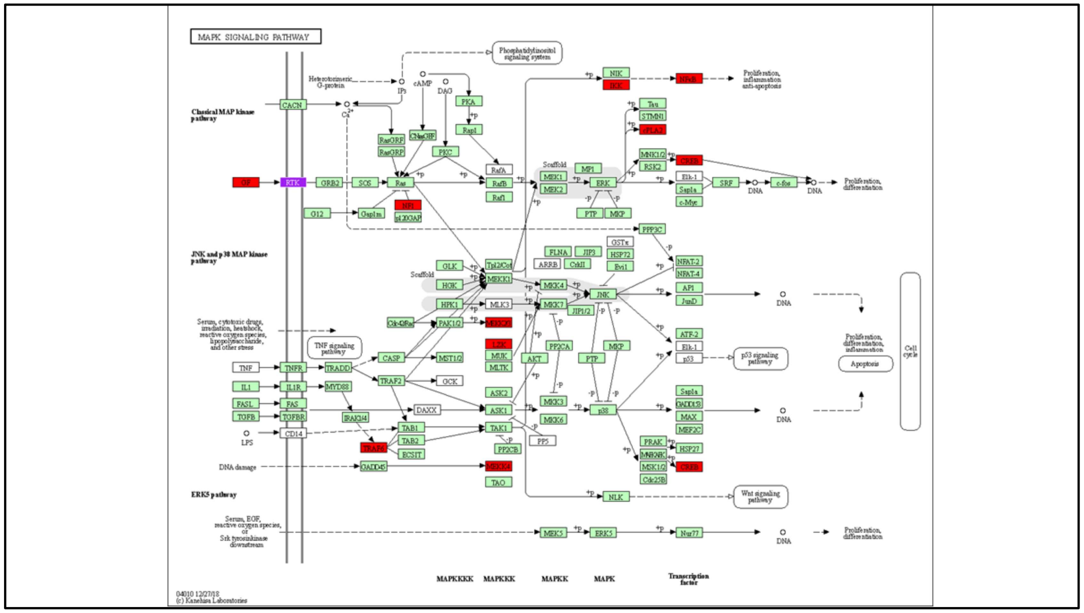

**Figure S34: The KEGG annotation of the MAPK signalling pathway.**

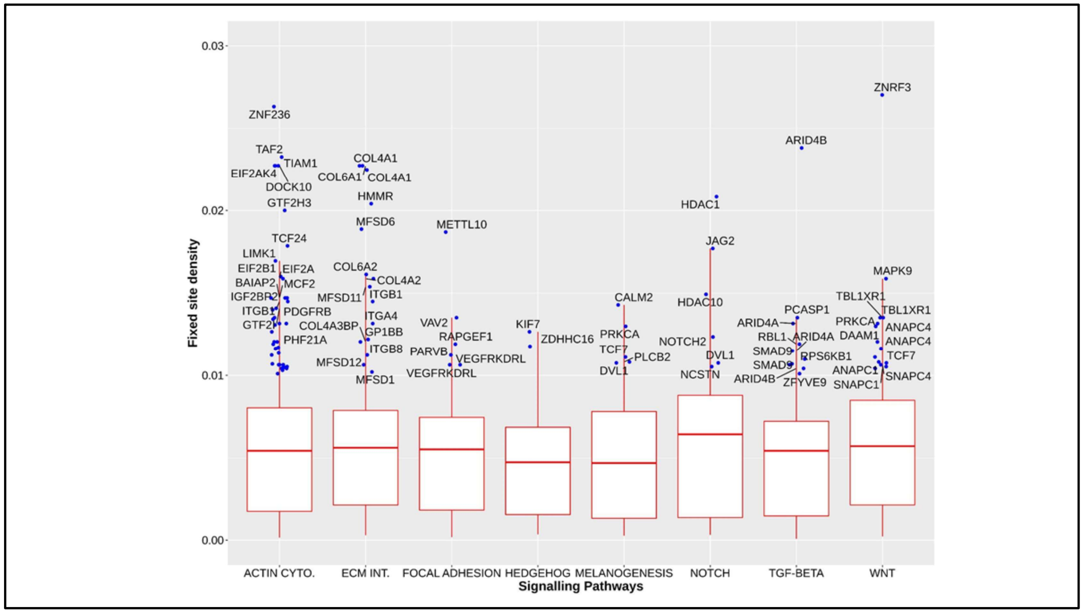

**Figure S35: The fixed site densities for the genes involved in various signalling pathways.**
Genes from several pathways (Actin cytoskeleton, Extracellular matrix interaction, Focal adhesion, Hedgehog, Melanogenesis, Notch, TGF-Beta and WNT) were screened to identify highly differentiated exons based on fixed site density. The fixed site density for each exon was calculated as the fraction of the number of fixed sites to the exon length. The exons of the genes showing fixed site density > 0.01 have been annotated on the plot.
